# Architectural singularities in wild Coffea (Baracoffea) species: integrated morphological perspectives for climate-resilient coffee cultivation

**DOI:** 10.1101/2024.12.18.628089

**Authors:** Rickarlos Bezandry, Romain Guyot, Hery Lisy Tiana Ranarijaona, Sylvie Sabatier, Marie Elodie Vavitsara, Artemis Anest

## Abstract

Global coffee production faces increasing threats from climate change, including rising temperatures, prolonged droughts, and spreads of diseases. Wild coffee species, notably those growthing in highly constrained environments in Madagascar, offer a critical genetic resource to address these challenges. Understanding adaptive traits allowing these species to establish into arid environments is essential to implement guid breeding strategies into create resilient coffee varieties. Here, we hypothesize that these wild Coffee species display unique traits compared to other Coffea species, reflecting adaptations to arid and seasonally dry environments. We used the architectural analysis to describe three Baracoffea species and compare them to known cultivated Coffee.

Field studies were conducted in two contrasting sites in Madagascar, focusing on three wild coffee species, natively growing into arid environments: Coffea ambongensis, C. bissetiae, and C. boinensis. Structural traits at the whole plant scale were measured across developmental stages using morphological and architectural analyses.

Our results suggest that unique traits characterize Barracoffea species, such as rhythmic growth, terminal flowering on short shoots, and species-specific developmental strategies. These findings highlight the architectural diversity of Baracoffea, identify potential key drivers of ecological adaptation and therefore highlights the potential of this group of species for breeding climate- resilient coffee varieties.

## Introduction

Coffee is a globally significant commodity, contributing over 200 billion dollars annually to the global economy (ICO, 2024; Samper et al., 2017). It is cultivated predominantly in tropical regions, providing livelihoods for more than 12.5 million smallholder farmers, and represents a cornerstone of economic stability for low and middle-income countries (ICO, 2024). Beyond its economic significance, coffee plays a vital cultural role, being an integral part of daily routines in many societies and a driver of social and economic interactions in both producing and consuming countries. However, coffee cultivation is increasingly threatened by climate change, which affects both yield and quality of production (Fig. 1).

**Fig. 1:**
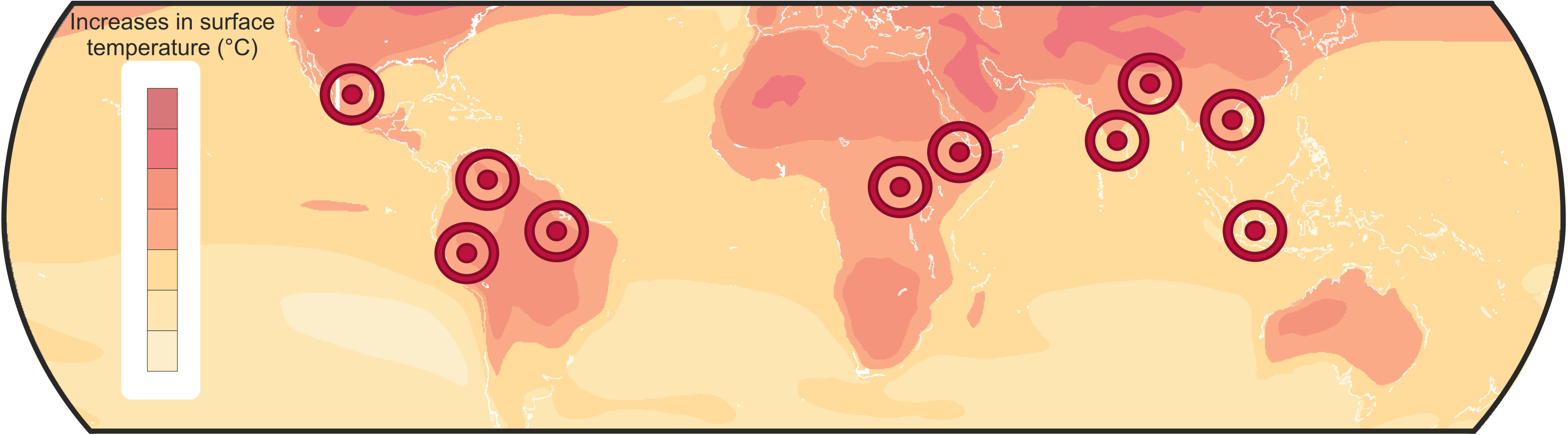
Map of the predictions in land surface temperature differences between now and 2050, modified from Di Lorenzo (2015) and with targets, indicating the top 10 countries in Coffee production (ICO, 2023).

Climate change constitutes a significant threat to coffee production (Bealu Girma, 2023; Bracken et al., 2021): rising temperatures, altered precipitation patterns, and extreme weather events impact coffee growth, development, and flavor (Bealu Girma, 2023). These changes are shifting suitable coffee-growing regions, with highland areas for Coffea arabica becoming too warm and lowland regions for Coffea canephora experiencing prolonged droughts and severe weather stress (Davis et al., 2012; Hamon et al., 2017). Projections suggest a 50% reduction in coffee-suitable land by 2050 globally, with Indonesia expected to lose over half of its Arabica-suitable area (Baker & Haggar, 2007, Davis et al., 2012; Ramadhillah & Masjud, 2024). Additionally, climate change is likely to increase the effect and presence of pest and disease (Davis et al., 2019, Bracken et al., 2021), further threatening coffee sustainability. Mitigation efforts, such as shade plantations and soil moisture control are commonly employed (Ramadhillah & Masjud, 2024), however, the widespread adoption of these strategies is limited by financial, environmental, and technical challenges, particularly for smallholder farmers. Addressing these barriers is essential to sustain coffee production and meet the rising demand (Bracken et al., 2021; Bracken et al., 2023).

While trying to address these challenges, coffee breeding programs remain constrained by the narrow genetic base of cultivated varieties. Coffea arabica, for instance, originated from a small genetic group of individuals (primarily in Ethiopia and Yemen) limiting its ability to adapt to new environmental stresses (Rakotomalala et al., 1992; Kang et al., 2014). Similarly, while Coffea canephora exhibits greater genetic diversity, its adaptive potential is still insufficient to face the magnitude of the effects of climate change. These genetic limitations emphasise the critical need for alternative sources of biodiverse material (Guyot et al., 2016, 2020, Bezandry et al., 2021a,b, 2023). Wild coffee species, particularly those endemic to Madagascar, represent a promising solution to these challenges. The Baracoffea group (Fig. 2), which includes nine species, is remarkably adapted to extreme conditions, such as arid climates and low fertility soils (Davis & Rakotonasolo, 2008; Bezandry et al., 2021). These species demonstrate a range of adaptations, including drought tolerance, high-temperature resilience, and pest resistance, making them valuable for breeding programs aimed at improving cultivated coffee (Davis et al., 2008). However, until now the potential of these species remains poorly studied, especially due to limited interest associated with low conservation efforts.

**Fig. 2:**
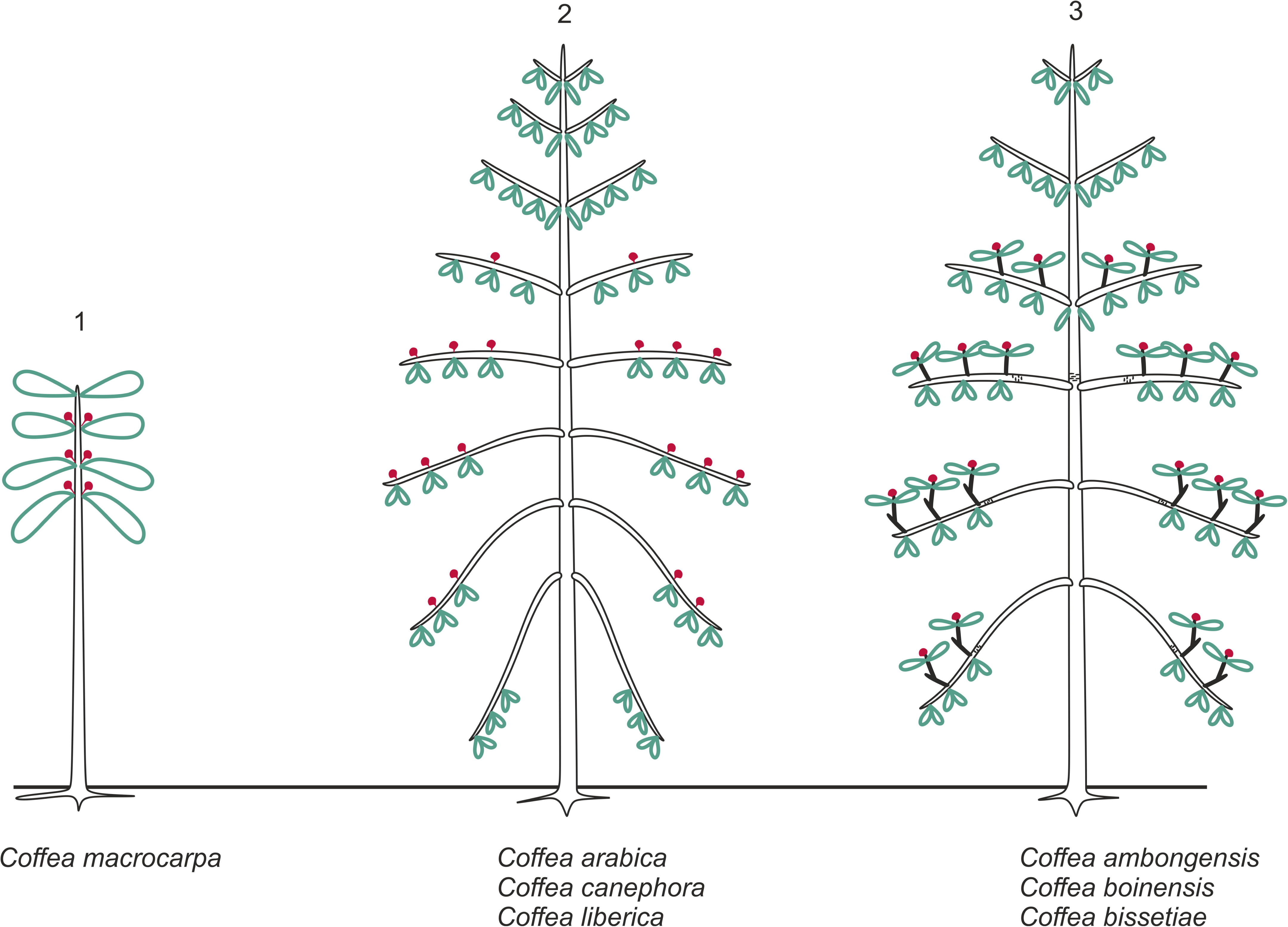
Three species of Baracoffea group, showing the whole plant (a) Coffea ambongensis Leroy ex Davis & Rakotonasolo, (b) Coffea bissetiae Davis & Rakotonasolo, and (c) Coffea boinensis Davis & Rakotonasolo; and detail of short shoots (c) C. ambongensis, (d) C. bissetiae and (e) C. boinensis. All photo credits: © Rickarlos Bezandry.

The conservation status of wild coffee species is indeed a growing concern. Over 70% of Madagascar’s native forest habitats have been lost due to deforestation and changes in land use, positioning many Baracoffea species at high risk of extinction (Myers et al., 2000; IUCN 2012,2023). Additionally, no comprehensive ex-situ conservation program has been created for these species, which are also not maintained in seed banks or botanical gardens (Ormsby & Kaplin, 2005; Davis & Rakotonasolo, 2021). Without immediate action, these genetic resources could be definitely lost.

Recent studies have demonstrated that specific structural traits can be crucial in plant adaptation to various environmental conditions, such as succulent organ establishment in arid environments (Arakaki et al., 2021). More specifically within these structural traits, plant architecture has been recently demonstrated to hold key characteristics to understand plant functional adaptations (Anest et al., 2024, Laurans et al., 2024).

Plant architectural analysis aims to describe the whole plant form and its establishment during plant development (Hallé et al., 1978; Oldeman, 1972; Edelin, 1974, 1984, 1993; Barthélémy, 1988, Barthélémy et al., 1989; Caraglio & Barthélémy, 1997), and decompose all this elements to connect them with ecological functions (Barthelemy & Caraglio, 2007; Laurans et al., 2024). Recent studies have demonstrated that plant architecture hold key functional traits in plant adaptation to aridity (Anest et al., 2021), or else defences to biotic disturbances (Charles-Dominique et al., 2017, Lefebvre et al., 2022, Anest et al, 2024). However, until now, only few attention has been addressed to plant architectural diversity in the Coffea genus, and how this could play a role in the ecological distribution of species and their tolerance to extreme conditions.

In this article, we aim to address these gaps by exploring the architectural and potentially adaptive traits of Baracoffea species that might explain their adaptation to extreme climatic conditions, and their potential for applications in coffee breeding programs. By understanding their ecological and architectural characteristics, this study aims to provide insights into the development of climate- resilient coffee varieties. These findings will also contribute to broader conservation efforts, making sure that wild coffee species can continue to serve as a crucial genetic reservoir for future generations.

Our hypotheses are that 1) the architectural traits of Baracoffea species are distinct to there relatives in Coffea genus and 2) reflect specific adaptations to their native environments, indicating their potential to enhance cultivated coffee’s resilience to climate-induced stresses (Davis et al., 2008; Bezandry et al., 2021). To test these hypotheses, we described and characterized the architecture of three Baracoffea species (Coffea ambongensis, Coffea bissetiae, and Coffea boinensis) to identify key architectural traits reflecting these adaptations.

## Materials and Methods

### Study sites and environmental context

The study was conducted in the northwestern Boeny Region of Madagascar, part of the western phytogeographical domain (Fig. S1,; Guillaumet & Koechlin, 1971). This region is characterized by seasonally dry tropical forests with predominantly deciduous vegetation. Water availability is the primary limiting factor for plant development in this area. The climate is tropical dry and hot, strongly influenced by the monsoon.

The study focused on two sites: Ankarafantsika National Park (PNA; longitude 46.57 to 47.28; latitude -16.00 to -16.33) and the Antsanitia forest (longitude 46.4 to 46.43; latitude -15.55 to - 15.58, both located in the Boeny Region. These sites were chosen for their contrasting ecological conditions and the presence of the Baracoffea species under study (Davis & Rakotonasolo, 2008; Bezandry et al., 2021).

Ankarafantsika (Fig. S2), spanning 136,513 hectares, comprises primary and secondary dry forests located 115 km south of Mahajanga. The park experiences two distinct seasons: a rainy season (November–April), with peak rainfall exceeding 500 mm in January, and a dry season (May– September), with rainfall as low as 5 mm in August. Average temperatures range from 22°C during the coolest months (June, July) to 29°C in October–December. Soils are predominantly sandy, derived from parent rock degradation, and prone to severe erosion. Vegetation includes seven natural habitat types, such as dry deciduous forests, riparian forests, and xerophilous thickets, hosting over 830 plant species (443 genera et 111 familles) with a 90% endemism rate.

The second site, Antsanitia forest (Fig. S3), represents degraded coastal forests and savannas. It covers 1,385 hectares and was designated as a national reforestation reserve in 1955.

Similar to PNA, Antsanitia has a six-month rainy season with peak rainfall above 400 mm in January, and a dry season with precipitation between 6–10 mm. Temperatures range from 24°C (July) to 28°C (October-December). Antsanitia’s vegetation includes coastal mangroves, savanna woodlands, and dry deciduous forests with drought-adapted species with around 68.6% being endemic (Rakotonandrasana et al., 2017). Soils range from sandy-clay to sandy-loam textures.

### Species selection and sampling strategy

This study focused on three species of the genus Coffea from the Baracoffea group (Davis & Rakotonasolo, 2008), within the Rubiaceae family: Coffea ambongensis Leroy ex Davis & Rakotonasolo, Coffea bissetiae Davis & Rakotonasolo, and Coffea boinensis Davis & Rakotonasolo (Fig. 2).

The Baracoffea species are shrubs or small trees, ranging from 1 to 6 meters tall, with a trunk diameter (at breast height) of up to 5 cm (Davis et al., 2005). The leaves are deciduous and bear domatia located at the intersection of the primary and secondary veins or occasionally between secondary and tertiary veins (Davis & Rakotonasolo, 2008).

The three studied species are naturally distributed across two sites in the Boeny region of western Madagascar: the Antsanitia forest and Ankarafantsika National Park. These species were selected for their accessibility, their shared general traits with other Baracoffea species (such as deciduous leaves and terminal flowering), and their distinct ecological habitats.

Coffea ambongensis is typically found in shaded understories, whereas Coffea bissetiae occupies intermediate habitats with moderate light exposure. Coffea boinensis mostly grows in open areas, making it an ideal candidate for studying adaptations to arid conditions (Bezandry et al., 2021).

1. C. ambongensis is found in the coastal and shrub forests of western Madagascar, particularly in Mahajanga Province (districts of Mahajanga II and Soalala). It grows on white sandy substrates at altitudes of 0–30 m and is likely to flower and fruit between October and March.
2. C. bissetiae is distributed along the western coast, in the districts of Marovoay, Mahajanga II, and Boriziny. It thrives in seasonally dry, deciduous forests on sandy or lateritic soils, at altitudes of 10– 240 m.
3. C. boinensis is restricted to Ankarafantsika National Park, in seasonally dry, deciduous forests on white sandy soils at altitudes of 170–210 m.

### Architectural and morphological measurements

Observations of individuals from the three Baracoffea species were conducted based on their theoretical stages of maturity, classified by branching order (Andrianasolo, 2012). Stage 1 represents unbranched individuals, Stage 2 includes individuals with A2 branches, Stage 3 features individuals with A3 shoots, and Stage 4 corresponds to adults with short shoots (A4).

For each developmental stage and species, a minimum of five individuals were observed, described, and measured, depending on availability. Selection criteria prioritized healthy individuals, free from visible damage, to ensure accurate representation of natural growth. Given the limited distribution of the three species across the two study sites (PNA and Antsanitia), sample sizes were occasionally restricted by availability and quality.

Between March and July 2021, 65 individuals were studied across the two sites. This included 25 individuals of C. boinensis, 26 of C. bissetiae, and 14 of C. ambongensis.

A total of 317 dry seeds were measured and weighed, comprising 222 seeds for C. boinensis, 69 for C. bissetiae, and 26 for C. ambongensis.

For foliar analysis, 960 dry leaves were measured, weighed, and scanned, distributed as follows: 300 leaves for C. boinensis, 360 for C. bissetiae, and 300 for C. ambongensis.

Given the conservation status of the studied species, non-destructive methods were employed, focusing on the observation and sketching of morphological traits. Efforts were made to examine as many trees as possible across different locations and developmental stages to generate a comprehensive representation of each species.

Observations were based on morphological concepts and criteria, following the architectural analysis protocols of Heuret et al. (2000) and Sabatier (1999). Analyses focused on identifying structural characteristics through morphological markers, including: Leaf scars defining internodes on stem segments, presence or absence of cataphylls, particularly at the apex; presence or absence of hypopodia at the base of shoots; bark features such as shape, size, and color; axis straightening; Architectural analysis was conducted following the Barthélémy and Caraglio (2007) framework.

Key descriptors used to characterize whole-plant architecture where included: 1) Phyllotaxy; 2) Growth direction and symmetry; 3) Meristematic activity and development; 4) Growth rhythmicity; 5) Spatial branching patterns, 6) Temporal branching patterns, 7) Sexuality position (see complete description of this traits in Barthelemy and Caraglio, 2007, and in Fig. S4).

For each developmental stage, at least five individuals were observed, sketched, and photographed. Observations were conducted in situ in forest environments, such as Ankarafantsika. In cases of taller trees, ropes were used to lower the trunks for detailed analysis.

All morphological traits were synthesized to identify and classify the different axis categories and architectural units specific to the studied species.

### Statistical and multivariate analysis

All analyses were performed using R version 4.3.1

Descriptive statistics were performed using the R software to calculate means, standard deviations, and variances for the variables studied. The Shapiro-Wilk test was employed to check the normality of the data distribution, and the Bartlett test was used to verify the homogeneity of variances.

Different statistical tests were applied based on the conditions of the data: Student’s t-test and ANOVA were used for parametric data. Mann-Whitney and Kruskal-Wallis tests were applied for non-parametric data.

Where significant differences between groups were detected, Tukey’s test (for parametric data) and Dunn’s test (for non-parametric data) were used for pairwise comparisons.

Spearman’s rank correlation test was employed to evaluate relationships between variables when data did not follow a normal distribution.

This analytical framework ensured accurate identification and interpretation of relationships between variables, providing a strong foundation for understanding species-specific traits.

## Results

### Architecture of Baracoffea species

Coffea ambongensis, Coffea bissetiae and Coffea boinensis (hereafter, Baracoffea group), as well as the cultivated species Coffea arabica and Coffea canephora, follow the Roux architectural model (according to Hallé et al., 1978), defined by a monopodial orthotropic trunk with continuous branching (hereafter referred to as the first axis category, C1) forming tiers of monopodial plagiotropic branches (second axis category, C2). According to the Roux model, sexuality is lateral, located on the trunk and branches. This differs in the species studied here, where sexuality is terminal and borne on short shoots (four axis category, C4).

The architectural model does not account for the presence of the most peripheral shoots (beyond the branches) and eclipse the information on terminal flowering on the short shoots in this variation of the Roux model (Fig. 3).

**Fig. 3:**
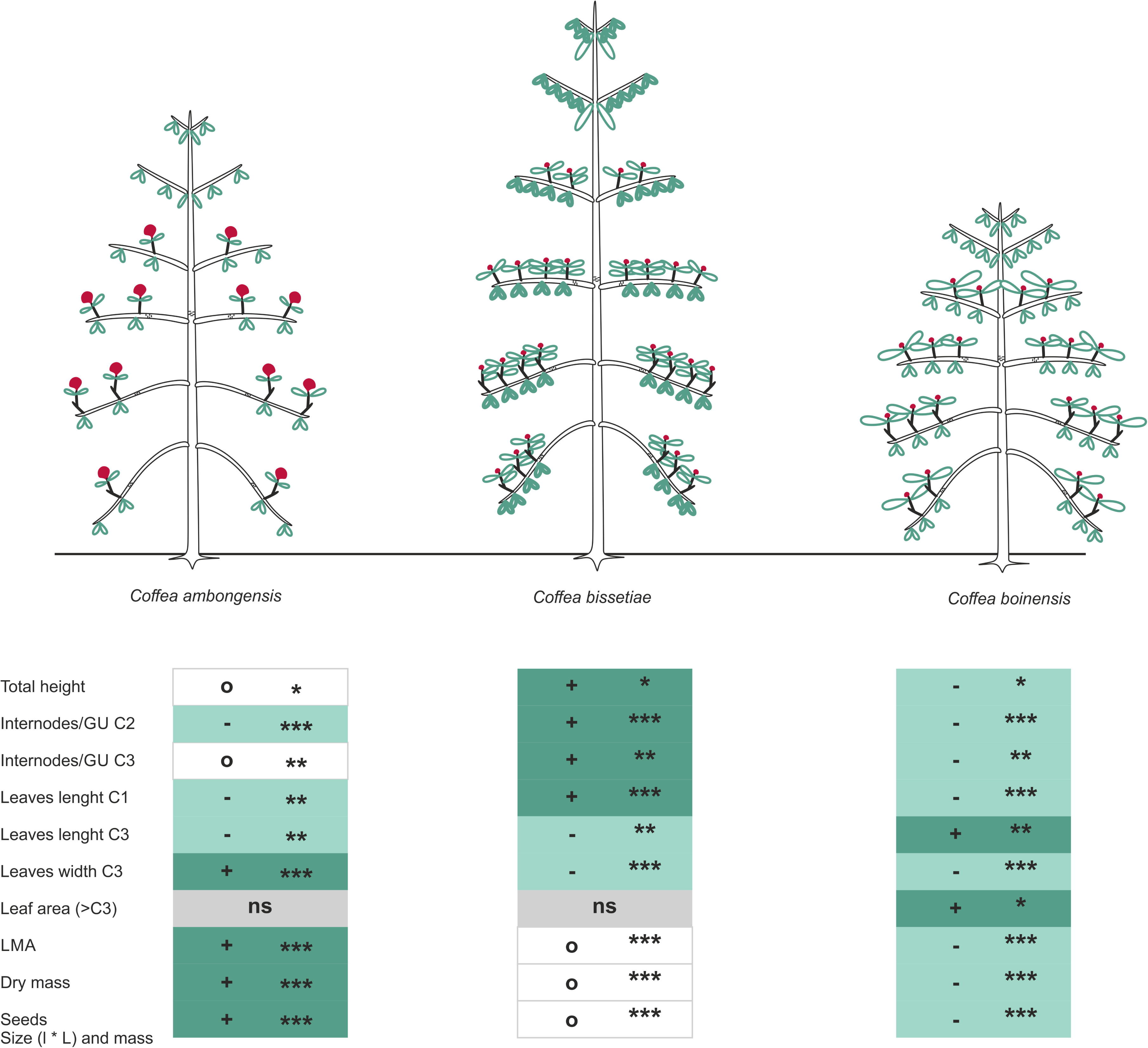
Three different architectures in genus Coffea. 1) Unbranched trunk with lateral flowering in Coffea macrocarpa, modified from Hallé (Hallé, 2019); 2) Typical Roux model in C. arabica, C. canephora et C. liberica modified from Hallé (Hallé, 2019), made of a monopodial orthotropic trunk with continuous branching forming tiers of monopodial plagiotropic branches which lateral flowering on the trunk and branches; 2) Variation of the Roux model in C. ambongensis, C. bissetiae and C. boinensis (this study), where sexuality is located in terminal position of short sympodial shoots, and growth is rhythmical. A detailed description of all tree species is available in Notes S1 to S3.

The three studied species differ from the other previously described Coffea species (Hallé, 2019) in their specific expression. Amphitonic branching (C3) appears continuously in a lateral position on the branches, themselve branched with shoots having very short internode (C4). Sexuality appears terminally, strictly on the ultimate axes (C4).

The main points of divergence with the other Coffea species (hereafter, Mascarocoffea group) is the structure of the architectural unit, with four categories in Baracoffea compared to three in Mascarocoffea. Additionally, the growth pattern, continuous in most Coffea species, is rhythmic in Baracoffeas (Fig. 4). Growth stops are identifiable by the setting up of a bud, protected by cataphylls (leaves transformed into scales). These cataphyll leave scars after the growth has restarted, delimiting the different growth units (abbreviated GU hereafter), i.e, uninterrupted portions of stem between two growth stops, representative of the annual growth. Finally, the position of sexuality, which is lateral on the branches in Mascarocoffea and terminal on the short shoots in Baracoffea. These characteristics are shared by all three Baracoffea species described, while they differ in some morphological aspects.

**Fig. 4:**
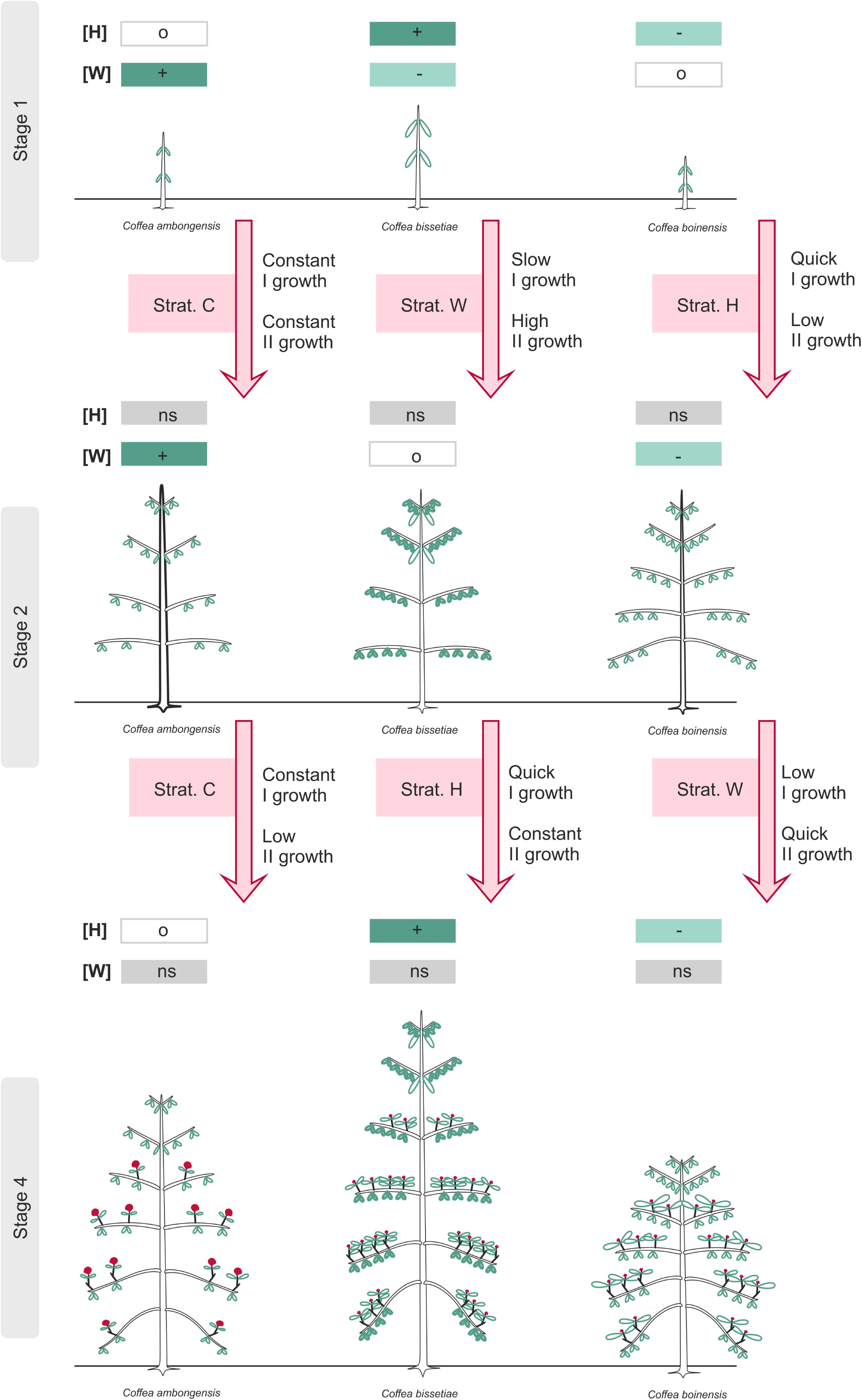
Scheme and summary of the morphological variations in the three Baracoffea species. + sign (associated with dark green) indicates significantly higher values, - sign (associated with light green) indicates significantly lower values and o sign (associated with white) indicates significantly different value and intermediate value, ns indicates no significant difference. All statistical tests outputs are in Supplementary Information. * Significant, ** Highly significant, *** Very highly significant, and NS Non-significant.

### Morphological variation in Baracoffea’s architecture

Where qualitatives characteristics in Baracoffea are similar for all three species, several traits different quantitatively (Fig. 4) Specifically, C. ambongensis is distinct by having only few internodes by growth units (mean 2.93 ± 0.59 cm, Table S1-S2, Fig. S5-S6) on the branches (C2), and shorter leaves (Table S4) on the trunk (C1; mean 5.25± 1.32 cm) and twigs (C3; mean 6.64 ± 1.80 cm). Leaves are however wider on twigs (C3; 3.89 ± 1.01 cm) than those other species, as well as having globally (all categories confunds) higher Leaf Mass per Area (LMA, Fig. S7; 0.026 ± 0.005 g/cm²) and dry mass (0.37 ± 0.06 g). Finally, the size (length and width; Fig. S8; 15.7 ± 1.4 and 11.35 ± 1.1 mm) and mass of seed (0.53 ± 0.1 g) are significantly the highest.

C. bissetiae is distinct by having the highest total height (Table S4; mean 358.73 ± 104.14 cm), the highest number of internodes by growth units (Table S2-S3, Fig. S5-S6) on branches and twigs (C2 and C3; respectively 4.35 ± 1.67 and 3.48 ± 1.22), and longer leaves (Table S4) on the trunk (C1; 7.58 ± 1.29 cm). Leaves are significantly smaller on twigs (C3; 7.19 ± 1.52 cm). Leaf Mass per Area (LMA, Fig. S7; 0.010 ± 0.005 g/cm²) and dry mass (0.07 ± 0.004 g) are also significantly the lowest . Finally, the size (length and width; respectively 6.33 ± 0.4 cm ; 5.68 ± 0.3 cm) and mass of seeds (Fig. S8; 0.06 ± 0.002 g) are significantly the lowest.

C. boinensis is distinct by having the lowest total height (Table S4; mean 190.61 ± 31.8 cm), and the lowest number of internodes by growth units (Table S1-S2, Fig. S5-S6) on twigs (C3; 2.53 ± 1.29). Leaves are significantly longer on twigs (C3; 9.21 ± 1.52 cm) and associated with higher leaf area on C3 (Fig. S9; 39.72 ± 1.2 cm²). Leaf Mass per Area (LMA, Fig. S7; 0.016 ± 0.005 g/cm²) and dry mass (0.26 ± 0.03 g/cm²) are also significantly lower than C. ambongensis and higher than C. bissetiae, as well as the size (length and width; respectively 14.32 ± 1.1 and 10.17 ± 0.9) and mass of seeds (Fig. S8; 0.29 ± 0.06 g).

These quantitative morphological variations result in contrasted shapes between the three species, and further results demonstrate that these differences are not established following the same pattern over species lifespan.

### Developmental strategies in Baracoffea

All three species show an ontogenetic path with similar developmental stages: stage 1 correspond to the seedling stage, when the plant is unbranched (only C1 present); stage 2 correspond to the stage plant setting up the first lateral branches (acquisition of C2); stage 3 correspond to the acquisition point of twig on branches (acquisition of C3); finally stage 4 is characterized by short shoot (C4) acquisition, expression of the sexuality and describes the adult individuals.

Where all the morphological events defining the stages appear in the same order in the three species lifespan, morphological quantitative variations shape each species in a distinct trajectory across stages (Fig. 5).

**Fig. 5:**
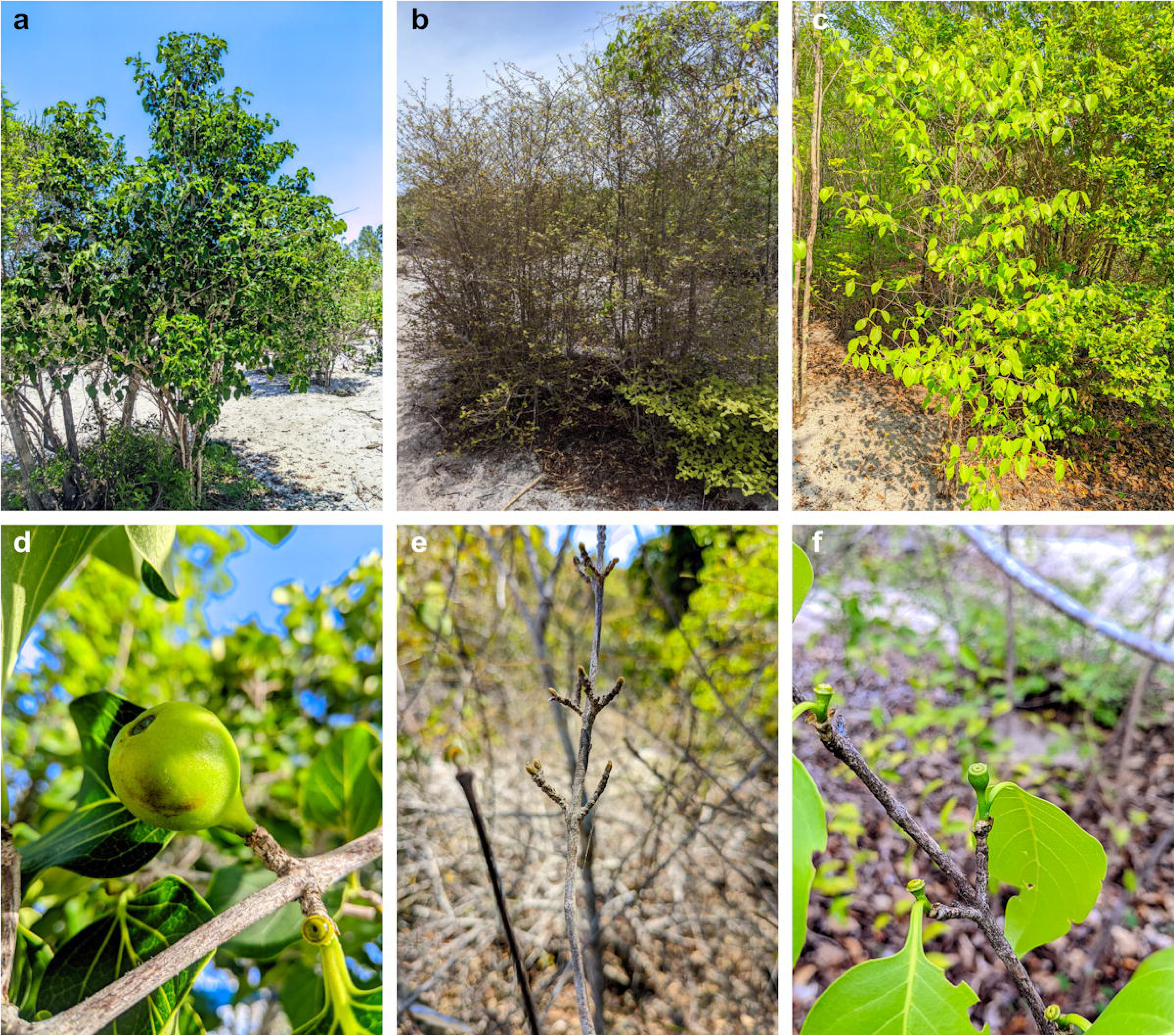
Scheme and summary of the morphological variations in the three Baracoffea species across different stages. Each line indicates a different stage. Here, stages 1, 2 and then 4 are represented. Stage 3 is not displayed as no significant differences occur between stage 2 and 3. [H] and [W] respect means “height” and “width” of the trunk. + sign indicates significantly higher values, - sign indicates significantly lower values and o sign indicates significantly different value and intermediate value, ns indicates no significant difference. All statistical tests outputs are in Supplementary Information. [Strat] indicates interpreted strategies of growth between stages where C means “Constant”, H means “Height” and W means “Width”. “I growth” means “primary growth” and “II growth” means “secondary growth”.

Specifically, during stage 1, C. bissetiae is significantly higher (14.10 ± 7.80 cm, Table S4, Fig. S10-S12) and has a thinner trunk diameter (0.21 ± 0.05 cm) than other species where C. boinensis is the smallest (7.05 ± 3.77 cm), and C. ambongensis the widest (0.22 ± 0.08 cm).

After reaching stage 2, all species show a similar high (average of 78.8 ± 26.16 cm). C. ambongensis remains the widest trunk diameter (1.65 ± 0.1 cm) where C. boinensis have the thinnest trunk diameter (0.48 ± 0.33 cm). This tandance is retained after reaching stage 3, with similar differences between species.

Finally, after reaching stage 4 and maturity (reproductive functions expressed), C. bissetiae is significantly higher than other species (358.73 ± 104.14 cm) and C. boinensis is the smallest (190.61 ± 31.8 cm). All species show a similar trunk diameter (average of 3.18 ± 0.88 cm).

Summarised at the species level, this indicates that C. ambongensis have a strategy of constant growth (Strategy C) during all stages of its developpement (Strategy C-C). Conversely, C. bissetiae display a not constant growth, with high secondary growth during first stages (Strategy W), then high primary growth (Strategy H, in fine Strategy W-H). Finally, C. boinensis show the opposite pattern, with high primary growth during first stages (Strategy H), then high primary growth (Strategy W, in fine Strategy H-W).

## Discussion

In this study we searched to identify structural traits of wild coffee species in Baracoffea group reflecting their adaptive potential to the extreme environment these species growth.

We identified unique sets of characters that are surprisingly contrasting with the morphology of Coffea described so far. More specifically, the rhythmic growth, marked by growth stops, a more complex architecture than common Coffea species and specific ontogenetic path are holding attention and will be further discussed.

### Rhythmicity of environment and of plants growth

Under water stress conditions, a key limiting factor for plant growth and survival (Ordoñez et al., 2010), two main strategies have been selected in plants: drought tolerance and drought avoidance. Drought tolerance involves mechanisms such as water storage (e.g., succulence) that maintain physiological processes while water availability is limited. In contrast, drought avoidance delays deficit of water by synchronizing physiological activities such as photosynthesis with favorable conditions (e.g., CAM photosynthesis, annual life cycle, deciduous foliage; Tomlinson et al., 2013). The morphological traits of Baracoffea species, compared to Mascarocoffea, suggest that the former has adopted the drought avoidance strategy (Reich and Borchert, 1984). In Baracoffea, this strategy manifests as deciduous foliage, unlike the evergreen leaves of Mascarocoffea, shedding leaves during the dry season as water potential decreases.

Additionally, drought avoidance in Baracoffea is associated with a stop of growth during unfavorable periods, resulting in rhythmic growth. This growth pattern alternates between a growth phase during favorable seasons, producing growth units (GUs), and a dormancy phase during dry seasons, marked by closer internodes, the presence of cataphylls, or reduced pith size. This differs from the continuous growth of Mascarocoffea.

These findings suggest that Baracoffea species structurally diverged from Mascarocoffea of Madagascar (Andrianasolo, 2012) and most Coffea species (Hallé et al., 1978; Okoma, 2019) through the selection of structural traits supporting drought avoidance, such as deciduous foliage and rhythmic growth. The seasonal leaf shedding in Baracoffea requires leaf renewal each favorable season, at the opposite of Mascarocoffea, which retains its foliage during the whole year. This renewal is facilitated by a unique structural trait: the presence of a new axis category, the short shoot.

### Complexity and functions in plant architecture

The very differentiated Baracoffea’s short shoots enable annual leaf production on short and resource efficient shoots (De Haldat et al., 2023). Some hypotheses also suggest that deciduous trees are often associated with short shoots to replace leaves without investing in new stem tissues (Dörken and Stützel, 2009). According to Anest et al. (2021), each new differentiated axis category has the potential to reinforce an existing function or enable a new one. In Baracoffea, in addition to low cost leaf regeneration (Dörken et al., 2010; Dörken and Stützel, 2009, 2012, De Haldat et al., 2023), the short shoots likely exponentially increased the number of fruiting sites, that are maintained over years through sympodial branching.

Despite these new traits, Baracoffea species retained continuous branching, a characteristic common among Coffea species (Andrianasolo, 2012; Hallé et al., 1978; Okoma & Sabatier, 2018). This may suggest that continuous branching either enhances resource exploitation during the rainy season, by increasing the number of short shoots and thus photosynthetic capacity, or is constrained by developmental limitations (Anest et al., 2021) that did not affect trait selection in arid environments.

The structural complexity of Baracoffea, with four specialized axis types, likely reflects a high degree of adaptation to their environment through functional specialization (Laurans, 2024; Anest, 2024). However, this complexity also implies greater sensitivity to environmental changes, as the evolution of new traits becomes more challenging (Anest et al., 2021). Complex architectures with more specialized axis categories are less suited to unstable or frequently disturbed ecosystems (Millan, 2016; Anest et al., 2021) but young stages are frequently less stress tolerant, and different developmental strategies might play a role in species survival.

### Ontogenetic trajectories, avoidance and resilience

While all the tree studied species are growing in the dry environment of Madagascar (Bezandry et al., 2023), contrasted ecological variables in their environment are likely witnessed by specific developmental trajectories in each species. In close environments, such as the Ankarafantsika sites where light availability is low, C. bissetiae develops a tall trunk, higher that other species, with numerous internodes, optimizing vertical growth to compete for light. Young stages display slow primary growth, suggesting different behavior than thus usually characterizing pioneer species that adopt rapid growth in height and diameter (Armani et et al., 2019) and suggesting this species is adapted to low light availability conditions (Fig. 6).

In contrast, in the more open environment (such as in Antsanitia), species develop with different strategies. C. ambongensis have a very linear growth suggesting that water is probably the single limiting factor in these systems. Conversely, C. boinensis display a very fast growing in height before slowing down after reaching the second and third stages of establishment. This strategy is common in fire dominated systems and might explain the necessity of fast growing to position vegetative parts and meristem quickly out of the fire reach (Hopkins et al., 2023, Fig. S13).

Our results emphasise the possibility of evolutionary divergence of the Baracoffea group (Rimlinger et al., 2020) and highlights the genetic potential of wild coffee species to diverse ecological conditions. These results reinforce the idea of the importance of structural traits in plant adaptability (i.e., adaptation and plasticity).

Implications for breeding: Threatened solutions

Our findings highlight the importance of understanding the architectural traits and associated adaptive strategies of Baracoffea species as a foundation for breeding programs. Comparing these results with morphological and quantitative traits of other Coffea species, particularly those growing in different environments (e.g., no limited water, other limiting factors), could provide valuable information. Additionally, expanding sample sizes to include more populations and diverse ecological conditions would enable further research to generate robust ecological and evolutionary analyses and integrate this knowledge into breeding programs. This approach could help anticipate the responses of coffee varieties to environmental constraints based on their structural traits and ecological factors that have selected these characteristics (Lauri, 2021; Maurin et al., 2023; Anest et al., 2024), offering predictive tools for breeding programs aiming to improve resilience in cultivated coffee trees.

However, this constitutes a significant challenge, as structural data on wild Coffea individuals are sparse, and these species are increasingly rare in their natural habitats. It is now critical to emphasize the importance of understanding the functioning and adaptations of these wild species, to highlight the value of preserving their genetic resources as a means to counteract the effects of climate change on coffee production (Gay, 2066; Jawo, 2022).

### Further requirements for breeding resistant and resilience Coffee varieties

Our study highlights the critical role of studying structural and functional adaptations of wild coffee species in addressing the challenges of climate change and improving the sustainability of coffee production. The possible adaptive traits observed in the Baracoffea group highlight their potential as a genetic resource for breeding climate-resilient coffee varieties. These results also emphasise the critical need for conservation efforts to protect these species from extinction.

By integrating this knowledge into breeding programs and conservation strategies, it is possible to secure the future of coffee as a global commodity. A multidisciplinary approach that combines genomics, agronomy, and ecology will be essential for achieving this goal, ensuring that coffee cultivation remains viable in the face of mounting environmental challenges.

## Acknowledgments

Rickalors Bezandry is sponsored by Rufford foundation grant n39692-1 and the French government grant awarded by the Service de coopération et d’action culturelle (SCAC) in Madagascar (N°104008W and N°122007W).

We are grateful to Madagascar National Parks (MNP) for allowing this research to be conducted.

This work has been made possible thanks to the Directorate of protected areas, renewable natural resources and ecosystems of Madagascar (Direction des aire protégées, des ressources naturelles renouvelables et des écosystèmes de Madagascar) research permit delivered N°270/24/SG/DAPRNE/SCBE.Re.

The authors gratefully acknowledge the support of the French Embassy in Madagascar and the Cooperation and Cultural Action Service (SCAC).

Rickarlos Bezandry has received the support of IRN LiStat (Statistics for Life Sciences and Ecology), the “Institut de Recherche pour le Développement” (IRD) and The French Agricultural Research Centre for International Development (CIRAD)

The authors thanks Frank RAKOTONASOLO for his help in the identification of Baracoffea species.

## Author Contribution

RB, SS et AA planned and designed the research, RB conducted data collection and processing. RB and AA analysed data. RB, HLTR, MEV, RG and AA wrote the manuscript.

## Data availability statement

The data associated with this article can be provided upon request to the authors.

## Supporting Information

Supporting Information associated to this article includes:

## Supporting tables

Table S1. Average number of internodes per growth unit and between species

Table S2. Average number of internodes per growth unit by type of axis within the same species

Table S3. Comparison of internodes and leaves for species at stage 4

Table S4. Description and comparison of species based on total height and basal diameter variables.

Table S5. Summary of the architectural unit of Coffea ambongensis

Table S6. Summary of the Architectural Unit of Coffea bissetiae Table S7. Recap of the architectural unit of Coffea boinensis Supporting figures

Fig. S1. Location of the two main study sites.

Fig. S2. Location and subdivision of the study zones within PNA.

Fig. S3. Location and subdivision of the study zones in Antsanitia.

Fig. S4. Some of the architectural descriptors commonly used in architectural analyses.

Fig. S5. Statistical comparison of the number of internodes per growth unit across species.

Fig. S6. Statistical comparison of the number of internodes per growth unit by axis type.

Fig. S7. Average leaf dry mass and LMA by species. Fig. S8. Average values of seed traits.

Fig. S9. Average leaf area (length x width, in cm²) by axis category.

Fig. S10. Height of the studied species across developmental stages.

Fig. S11. : Height and basal diameter from seedling to adult stage.

Fig. S12. Average internode length and leaf length at developmental stage 4

Fig. S13. Substrate of the Baracoffea environments

Fig. S14. Phyllotaxy in C. ambongensis and branched system with monopodial development in C. ambongensis.

Fig. S15. Rhythmic growth and morphological markers in C. ambongensis. Fig. S16. Immediate and delayed branching in C. ambongensis.

Fig. S17. Terminal Sexuality on Short Shoots in Coffea ambongensis.

Fig. S18: Opposite-Decussate Phyllotaxy in Coffea bissetiae.

Fig. S19. Monopodial branched system in Coffea bissetiae and rhythmic growth and morphological marker of growth.

Fig. S20. Immediate branching in Coffea bissetiae.

Fig. S21. Terminal sexuality on short shoots in Coffea bissetiae.

Fig. S22. Opposite-decussate phyllotaxy in Coffea boinensis.

Fig. S23. Monopodial branching system, rhythmic growth and morphological markers in Coffea boinensis.

Fig. S24. Immediate branching in Coffea boinensis.

Fig. S25. Terminal sexuality in Coffea boinensis.

## Supporting notes

Notes. S1. Architectural and morphological description of Coffea ambongensis. Notes. S1. Architectural and morphological description of Coffea bissetiae.

Notes. S3. Architectural and morphological description of Coffea boinensis.

## References

Andrianasolo, D. N. (2012). Génétique des populations et modèles d’architecture et de production végétale: application à la préservation des ressources génétiques des Mascarocoffea (Doctoral dissertation, Montpellier 2).

Anest, A., T. Charles-Dominique, O. Maurin, M. Millan, C. Edelin et K. W. Tomlinson (2021). Evolving the structure: climatic and developmental constraints on the evolution of plant architecture. A case study in Euphorbia New Phytologist, vol. 231 : 1278lll1295.

Anest, A., Bouchenak-Khelladi, Y., Charles-Dominique, T., Forest, F., Caraglio, Y., Hempson, G. P., … & Tomlinson, K. W. (2024). Blocking then stinging as a case of two-step evolution of defensive cage architectures in herbivore-driven ecosystems. Nature Plants, 10, 587–597.

Arakaki, M., Christin, P. A., Nyffeler, R., Lendel, A., Eggli, U., Ogburn, R. M., … & Edwards, E. J. (2011). Contemporaneous and recent radiations of the world’s major succulent plant lineages. Proceedings of the National Academy of Sciences, 108(20), 8379–8384.

Armani, M., Charles-Dominique, T., Barton, K. E., & Tomlinson, K. W. (2019). Developmental constraints and resource environment shape early emergence and investment in spines in saplings. Annals of Botany, 124(7), 1133–1142.

Baker, P. et J. Haggar (2007). Global Warming: the impact on global coffee. In SCAA conference handout. Long Beach May 2007, USA : 14p.

Barthélémy, D. (1988). Architecture and sexuality in some tropical plants: the concept of automatic flowering, Architecture et sexualité chez quelques plantes tropicaleslll: le concept de floraison automatique. Université des Sciences et Techniques du Languedoc, Montpellier, France. 262 pages. https://hal.science/tel-03817694

Barthélémy, D., C. Edelin et F. Hallé (1989). Architectural concepts for tropical trees In H. Balslev et I. Nielsen, Éds. Tropical Forests: Botanical Dynamics, Speciation & Diversity. Academic Press : 89–100.

Barthélémy, D. et Y. Caraglio (2007). Plant Architecture: A Dynamic, Multilevel and Comprehensive Approach to Plant Form, Structure and Ontogeny. Annals of Botany, vol. 99 : 375lll407. 10.1093/aob/mcl260

Bezandry, R., S. Sabatier, R. Guyot et M. Vavitsara (2021a). Architecture et variabilité interspécifique de trois espèces de Baracoffealll: Coffea boinensis, Coffea bissetiae et Coffea ambongensis. Revue des Sciences, de Technologies et de l’Environnement. Édition spéciale, Université d’été 3ème édition Mahajanga, novembre 2021, vol. Volume 5 : 8lll16.

Bezandry, R., Vatvitsara, M. E., Rakotonasolo, F., Guyot, R., & Sabatier, S. A. (2021b). Studies of the Baracoffea: Malagasy coffee trees growing on the West Coast of Madagascar. ASIC. Bezandry, R., Dupeyron, M., Gonzalez-Garcia, L. N., Anest, A., Hamon, P., Ranarijaona, H. L. T.,

Bezandry, R., Vatvitsara, M. E., Rakotonasolo, F., Guyot, R. & Guyot, R. (2024). The evolutionary history of three Baracoffea species from western Madagascar revealed by chloroplast and nuclear genomes. Plos one, 19, e0296362.

Bracken, P., Burgess, P. J., & Girkin, N. T. (2021). Enhancing the climate resilience of coffee production. agriRxiv, (2019), 20210490350.

Bracken, P., Burgess, P. J., & Girkin, N. T. (2023). Opportunities for enhancing the climate resilience of coffee production through improved crop, soil and water management. Agroecology and Sustainable Food Systems, 47(8), 1125–1157.

Caraglio, Y. et D. Barthélémy (1997). Revue critique des termes relatifs à la croissance et à la ramification des tiges des végétaux vasculaires. Arbres et Sciences, vol. 13 : 1lll61.

Charles-Dominique, T., Barczi, J. F., Le Roux, E., & Chamaillé-Jammes, S. (2017). The architectural design of trees protects them against large herbivores. Functional Ecology, 31, 1710–1717.

Davis, A. P. et F. Rakotonasolo (2008). A taxonomic revision of the baracoffea alliance: nine remarkable Coffea species from western Madagascar Botanical Journal of the Linnean Society, vol. 158 : 355lll390. 10.1111/j.1095-8339.2008.00936.x.

Davis, A. P., F. Rakotonasolo et P. De Block (2010). Coffea toshii sp. nov. (Rubiaceae) from Madagascar. Nordic Journal of Botany, vol. 28 : 134lll136. 10.1111/j.1756-1051.2010.00710.x.

Davis, A. P. (2011). Psilanthus mannii, the type species of Psilanthus, transferred to Coffea. Nordic Journal of Botany, vol. 29 : 471lll472.

Davis, A. P., T. W. Gole, S. Baena et J. Moat (2012). The Impact of Climate Change on Indigenous Arabica Coffee (Coffea arabica): Predicting Future Trends and Identifying Priorities. PLoS ONE, vol. 7 (11) : e47981. 10.1371/journal.pone.0047981.

Davis, A. P., H. Chadburn, J. Moat, R. O’Sullivan, S. Hargreaves et E. Nic Lughadha (2019). High extinction risk for wild coffee species and implications for coffee sector sustainability. Science Advances, vol. 5 (1) : eaav3473. 10.1126/sciadv.aav3473.

Davis, A. P. et F. Rakotonasolo (2021). Six new species of coffee (Coffea) from northern Madagascar. Kew Bulletin, vol. 76 : 497lll511. 10.1007/s12225-021-09952-5.

De Haldat Du Lys, A., Millan, M., Barczi, J. F., Caraglio, Y., Midgley, G. F., & Charles-Dominique, T. (2023). If self-shading is so bad, why is there so much? Short shoots reconcile costs and benefits. New Phytologist, 237(5), 1684–1695.

Di Lorenzo, E. (2015). The future of coastal ocean upwelling. Nature, 518(7539), 310–311.

Dörken, V. M. et T. Stützel (2009). The adaptive value of shoot differentiation in deciduous trees and its evolutionary relevance. Boletín de la Sociedad Argentina de Botánica, vol. 44 : 421lll439.

Dörken, V. M., G. Stephan et T. Stützel (2010). Morphology and anatomy of anomalous short shoots in Pinus (Pinaceae) and their evolutionary meaning. Feddes Repertorium, vol. 121 : 133lll155. 10.1002/fedr.201000006.

Dörken, V. M. et T. Stützel (2012). Morphology, anatomy and vasculature of leaves in Pinus (Pinaceae) and its evolutionary meaning. Flora - Morphology, Distribution, Functional Ecology of Plants, vol. 207 : 57lll62. 10.1016/j.flora.2011.10.004.

Edelin, C. (1977). Images de l’architecture des conifères. Université de Montpellier 2. Thèse de doctorat de 3e Cycle. 255 pages.

Edelin, C. (1984). L’architecture monopodiale : l’exemple de quelques arbres d’Asie tropicale. Université de Montpellier 2. These de doctorat.

Edelin, C. (1993). Aspects morphologiques de la croissance rythmique chez les arbres tropicaux. Compte Rendu du Séminaire du groupe d’Étude de l’Arbre : le rythme de croissance, base de l’organisation temporelle de l’arbre. Angers, 13lll23.

Eilam, T., Y. Anikster, E. Millet, J. Manisterski, O. Sagi-Assif et M. Feldman (2007). Genome size and genome evolution in diploid Triticeae species. Genome, vol. 50 : 1029lll1037. 10.1139/G07-083.

Gay, C., F. Estrada, C. Conde, H. Eakin et L. Villers (2006). Potential Impacts of Climate Change on Agriculture: A Case of Study of Coffee Production in Veracruz, Mexico. Climatic Change, vol. 79 : 259lll288. 10.1007/s10584-006-9066-x.

Gokavi, N. et M. Kishor (2020). Impact of climate change on coffee production: An overview 9.

Guillaumet, J. L., & Koechlin, J. (1971). Contribution à la définition des types de végétation dans les régions tropicales (exemple de Madagascar). Candollea, 26(2), 263-277.

Guyot, R., T. Darré, M. Dupeyron, A. de Kochko, S. Hamon, E. Couturon, D. Crouzillat, M. Rigoreau, J.J. Rakotomalala, N. E. Raharimalala, S. D. Akaffou et P. Hamon (2016). Partial sequencing reveals the transposable element composition of Coffea genomes and provides evidence for distinct evolutionary stories. Molecular Genetics and Genomics, vol. 291 : 1979lll1990. 10.1007/s00438-016-1235-7.

Guyot, R., P. Hamon, E. Couturon, N. Raharimalala, J.J. Rakotomalala, S. Lakkanna, S. Sabatier, A. Affouard et P. Bonnet (2020). WCSdb: a database of wild Coffea species. DatabaseflJ: The Journal of Biological Databases and Curation, vol. 2020 . 10.1093/database/baaa069.

Girma, B. (2023). Climate Change and Coffee Quality: Challenges and Strategies for a Sustainable Future. Advances in Bioscience and Bioengineering, 11(2), 27.

Hallé, F., P. B. Tomlinson et M. H. Zimmermann (1978). Architectural variation at the specific level in tropical trees. Tropical trees as living systems, 209lll221.

Hamon, P., C. E. Grover, A. P. Davis, J.J. Rakotomalala, N. Raharimalala, V. A. Albert, H. L. Sreenath, P. Stoffelen, S. E. Mitchell, E. Couturon, S. Hamon, A. de Kochko, D. Crouzillat, M. Rigoreau, U. Sumirat, S. Akaffou et R. Guyot (2017). Genotyping-by-sequencing provides the first well-resolved phylogeny for coffee (Coffea) and insights into the evolution of caffeine content in its species. Molecular Phylogenetics and Evolution, vol. 109 351lll361. 10.1016/j.ympev.2017.02.009.

Heuret, P. (2002). Analysis and modelling of sequences of botanical events: understanding the regularity of expression of the growth, branching and flowering processes, Analyse et modélisation de séquences d’évènements botaniques: applications à la compréhension de la régularité d’expression des processus de croissance, de ramification et de floraison. Université Henri Poincaré, Nancy I. Thèse de doctorat. 145 ICO (2019a). Coffe Developpement Report 2019lll: Growing for Prosperity - Economic Viability as the Catalyst for a Sustainable Coffee Sector. http://www.ico.org/documents/cy2021-22/coffee-development-report-2019.pdf.

Hopkins, J. R., Huffman, J. M., Jones, N. J., Platt, W. J., & Sikes, B. A. (2023). Pyrophilic plants respond to postfire soil conditions in a frequently burned longleaf pine savanna. The American Naturalist, 201(3), 389–403.

International Coffee Organization, ICO (2023). Coffee report and outlook (CRO). Available at: https://icocoffee.org/documents/cy2023-24/Coffee_Report_and_Outlook_December_2023_ICO.pdf (Accessed December 7, 2024).

Jawo, T. O., D. Kyereh et B. Lojka (2022). The impact of climate change on coffee production of small farmers and their adaptation strategies: a review. Climate and Development, vol. 0 : 1lll17. 10.1080/17565529.2022.2057906.

Kang, M., J. Tao, J. Wang, C. Ren, Q. Qi, Q. Xiang et H. Huang (2014). Adaptive and nonadaptive genome size evolution in Karst endemic flora of China. New Phytologist, vol. 202 : 1371lll1381. 10.1111/nph.12726.

Laurans, M., Munoz, F., Charles-Dominique, T., Heuret, P., Fortunel, C., Isnard, S., … & Violle, C. (2024). Why incorporate plant architecture into trait-based ecology?. Trends in Ecology & Evolution.

Lauri, P. É. (2021). Tree architecture and functioning facing multispecies environments: We have gone only halfway in fruit-trees. American Journal of Botany, 108.

Lefebvre, T., Charles-Dominique, T., & Tomlinson, K. W. (2022). Trunk spines of trees: a physical defence against bark removal and climbing by mammals?. Annals of Botany, 129(5), 541–554.

Maurin, O., A. P. Davis, M. Chester, E. F. Mvungi, Y. Jaufeerally-Fakim et M. F. Fay (2007). Towards a Phylogeny for Coffea (Rubiaceae): Identifying Well-supported Lineages Based on Nuclear and Plastid DNA Sequences. Annals of Botany, vol. 100 : 1565lll1583. 10.1093/aob/mcm257.

Maurin, O., Anest, A., Forest, F., Turner, I., Barrett, R. L., Cowan, R. C., … & Charles-Dominique, T. (2023). Drift in the tropics: Phylogenetics and biogeographical patterns in Combretaceae. Global Ecology and Biogeography, 32(10), 1790–1802.

Myers, N., R. A. Mittermeier, C. G. Mittermeier, G. A. Da Fonseca et J. Kent (2000). Biodiversity hotspots for conservation priorities. Nature, vol. 403 : 853lll858.

Okoma, M. P., & Sabatier, S. A. (2018). Modélisation des paramètres de croissance et de développement chez différentes espèces de caféiers et Côte d’Ivoire. CIRAD.

Oldeman, R. A. (1972). L’architecture de la forêt guyanaise. Université de Montpellier 2, France. Thèse de doctorat, 247 pages.

Ordoñez, J. C., Van Bodegom, P. M., Witte, J. P. M., Bartholomeus, R. P., Van Dobben, H. F., & Aerts, R. (2010). Leaf habit and woodiness regulate different leaf economy traits at a given nutrient supply. Ecology, 91(11), 3218–3228.

Ormsby, A. et B. A. Kaplin (2005). A framework for understanding community resident perceptions of Masoala National Park, Madagascar. Environmental Conservation, vol. 32 : 156lll164. 10.1017/S0376892905002146.

Ramadhillah, B., & Masjud, Y. I. (2024). Climate change impacts on coffee production in Indonesia: A review. Journal of Critical Ecology, 1(1), 1–7.

Rakotomalala, J. J. (1992). Diversité biochimique des caféiers : analyse des acides hydroxycinnamiques, bases puriques et diterpènes glycosidiques. Particularités des caféiers sauvages de la région malgache (Mascarocoffea Chev.). Sciences et Techniques du Languedoc, Université de Montpellier 2 (Collection TDM - ORSTOM). Thèse de Doctorat, 217 pages.

Rakotonandrasana, S., Rakotondrafara, A., Rakotondrajaona, R., Rasamison, V., & Ratsimbason, M. (2017). Plantes médicinales des formations végétales de la baie de Rigny- Antsiranana à Madagascar. BOIS & FORETS DES TROPIQUES, 331, 55–65.

Reich, P. B., & Borchert, R. (1984). Water stress and tree phenology in a tropical dry forest in the lowlands of Costa Rica. The Journal of Ecology, 61-74.

Rimlinger, A., Raharimalala, N., Letort, V., Rakotomalala, J. J., Crouzillat, D., Guyot, R., … & Sabatier, S. (2020). Phenotypic diversity assessment within a major ex situ collection of wild endemic coffees in Madagascar. Annals of Botany, 126(5), 849–863.

Sabatier, S. (1999). Variabilité morphologique et architecture de deux espèces de noyers: Juglans regia L., Juglans nigra L. et de deux noyers hybrides interspecifiques (Doctoral dissertation, Université Montpellier II-Sciences et Techniques du Languedoc).

Samper, L. F., D. Giovannucci et L. M. Vieira (2017). The powerful role of intangibles in the coffee value chain. WIPO, vol. 39.

Tomlinson, K. W., Poorter, L., Sterck, F. J., Borghetti, F., Ward, D., de Bie, S., & van Langevelde, F. (2013). Leaf adaptations of evergreen and deciduous trees of semi-arid and humid savannas on three continents. Journal of Ecology, 101, 430–440.

IUCN (2012). Catégories et Critères de la Liste rouge de l’UICN : Version 3.1. Deuxième édition. Gland, Suisse et Cambridge, Royaume-Uni : UICN.

IUCN (2023). The IUCN Red List of Threatened Species IUCN Red List of Threatened Species. https://www.iucnredlist.org/fr.

