## Supplementary material for "Architectural singularities in wild Coffea (Baracoffea) species: integrated morphological perspectives for climate-resilient coffee cultivation"

### Supporting Information

The present supporting Information Includes:

### Supporting tables

**Table S1.** Average number of internodes per growth unit and between species

The number of internodes per growth unit varies depending on the species and the type of axis.

**Table S2.** Average number of internodes per growth unit by type of axis within the same species

The number of internodes per growth unit varies depending on the species and the type of axis considered.

**Table S3.** Comparison of internodes and leaves for species at stage 4

The studied traits include: leaf length and width, internode length (I.N. Length) and diameter (I.N. Diam.), number of internodes (I.N. nb).

**Table S4.** Description and comparison of species based on total height and basal diameter variables.

Total height and basal diameter vary according to the developmental stage of individuals across all species.

**Table S5.** Summary of the architectural unit of *Coffea ambongensis*

**Table S6.** Summary of the Architectural Unit of *Coffea bissetiae*

**Table S7.** Recap of the architectural unit of *Coffea boinensis*

### Supporting figures

**Fig. S1.** Location of the two main study sites.

**Fig. S2.** Location and subdivision of the study zones within PNA.

**Fig. S3.** Location and subdivision of the study zones in Antsanitia.

**Fig. S4.** Some of the architectural descriptors commonly used in architectural analyses.

**Fig. S5.** Statistical comparison of the number of internodes per growth unit across species.

Graphical representations of the Kruskal-Wallis test results for the number of internodes per growth unit (GU) for the trunk, branches, and shoots, respectively.

**Fig. S6.** Statistical comparison of the number of internodes per growth unit by axis type.

Graphical representations of the Kruskal-Wallis test.

**Fig. S7.** Average leaf dry mass and LMA by species.

**Fig. S8.** Average values of seed traits.

**Fig. S9.** Average leaf area (length x width, in $cm^2$) by axis category.

**Fig. S10.** Height of the studied species across developmental stages.

**Fig. S11. :** Height and basal diameter from seedling to adult stage.

**Fig. S12**. Average internode length and leaf length at developmental stage 4

**Fig. S13.** Substrate of the Baracoffea environments

**Fig. S14.** Phyllotaxy in *C. ambongensis* and branched system with monopodial development in *C. ambongensis*.

**Fig. S15.** Rhythmic growth and morphological markers in *C. ambongensis*.

**Fig. S16.** Immediate and delayed branching in *C. ambongensis.*

**Fig. S17.** Terminal Sexuality on Short Shoots in *Coffea ambongensis*.

**Fig. S18:** Opposite-Decussate Phyllotaxy in *Coffea bissetiae*.

**Fig. S19.** Monopodial branched system in *Coffea bissetiae* and rhythmic growth and morphological marker of growth.

**Fig. S20.** Immediate branching in *Coffea bissetiae*.

**Fig. S21.** Terminal sexuality on short shoots in *Coffea bissetiae*.

**Fig. S22.** Opposite-decussate phyllotaxy in *Coffea boinensis.*

**Fig. S23.** Monopodial branching system, rhythmic growth and morphological markers in *Coffea boinensis*.

**Fig. S24.** Immediate branching in *Coffea boinensis*.

**Fig. S25.** Terminal sexuality in *Coffea boinensis*.

### Supporting notes

**Notes. S1.** Architectural and morphological description of *Coffea ambongensis*.

**Notes. S1.** Architectural and morphological description of *Coffea bissetiae*.

**Notes. S3.** Architectural and morphological description of *Coffea boinensis*.

**Supplementary tables**

**Table S1. Average number of internodes per growth unit and between species**

The number of internodes per growth unit varies depending on the species and the type of axis considered in trees at developmental stage 4 (adult individuals). For *C. ambongensis* and *C.* boinensis, the number of internodes per growth unit varies significantly between the main axis and the peripheral axes (p-value < 0.01). The number of internodes per growth unit progressively decreases from the trunk to the most differentiated axes in all three species, measuring 4.82 (±2.51), 2.93 (±1.56), and 2.90 (±1.25) in the trunk (C1), branches (C2), and shoots (C3), respectively.

Significance levels for the Kruskal-Wallis test: * Significant, ** Highly significant, *** Very highly significant, and NS Non-significant. AMB: *C. ambongensis*, BIS: *C. bissetiae*, BOI: *C. boinensis*.

| **Axes categories** | **AMB** | **BIS** | **BOI** | **Pr>F** | **Tests (axes)** |
|---|---|---|---|---|---|
| **Trunk (C1)** | 4.82±2.51 a | 3.91±1.81 a | 4.50±2.72 a | $0.26^{NS}$ | Kruskal-Wallis |
| **Branches (C2)** | 2.93±1.59 b | 4.35±1.67 a | 3.09±1.78 b | **$7.00e^{-5}$ ***** | Kruskal-Wallis |
| **Twigs (C3)** | 2.90±1.25 ab | 3.48±1.22 a | 2.53±1.29 b | **0.0031 **** | Kurskal-Wallis |

**Table S2. Average number of internodes per growth unit by type of axis within the same species**

The number of internodes per growth unit varies depending on the species and the type of axis considered in trees at developmental stage 4 (adult individuals). Along the main axis, the number of internodes shows minor variation, ranging from 3.91 (±1.81) to 4.82 (±2.51) across the three studied species. Statistically significant variations are observed in the branches and shoots among the three species (p-value < 0.01). C. bissetiae has the highest number of internodes per growth unit in both branches (4.35 ± 1.67) and shoots (3.48 ± 1.22).

Significance levels for the Kruskal-Wallis test: * Significant, ** Highly significant, *** Very highly significant, and NS Non-significant. AMB: *C. ambongensis*, BIS: *C. bissetiae*, BOI: *C. boinensis*.

| Species | Trunk (C1) | Branches (C2) | Twigs (C3) | Pr>F | Tests |
|---|---|---|---|---|---|
| **AMB** | 4.82±2.51 a | 2.93±1.56 b | 2.90±1.25 b | 0.00017*** | Kruskall-Wallis |
| **BIS** | 3.91±1.81 a | 4.35±1.67 a | 3.48±1.22 a | 0.13 $^{NS}$ | Kruskall-Wallis |
| **BOI** | 4.50±2.72 a | 3.09±1.78 ab | 2.53±1.29 b | 0.0022 ** | Kruskall-Wallis |

**Table S3. Comparison of internodes and leaves for species at stage 4**

The studied traits include: leaf length and width, internode length (I.N. Length) and diameter (I.N. Diam.), number of internodes (I.N. nb).

Significance levels for the Kruskal-Wallis test: * Significant, ** Highly significant, *** Very highly significant, and NS Non-significant.

AMB: C. *ambongensis*, BIS: C. *bissetiae*, BOI: C. *boinensis*.

| Species | I.N. Length (cm) | I.N. Diam. (cm) | I.N. nb | Leaf length (cm) | Leaf width (cm) |
|---|---|---|---|---|---|
| BOI C1 | 8.66±5.50 a | 1.20±0.86 b | 22±4 b | 8.95±1.29 a | 3.77±0.74 a |
| BIS C1 | 7.47±6.57 b | 1.45±0.80 a | 48±15 a | 7.58±1.29 a | 3.12±0.48 a |
| AMB C1 | 6.55±6.33 c | 1.45±1.11 ab | 38±7.6 a | 5.25±1.32 b | 3.88±1.11 a |
| **Pr>F** | 8.947e-09*** | 0.005** | 0.0009*** | 0.000188*** | 0.062 NS |
| BOI C2 | 5.66±3.50 a | 0.36±0.12 b | 8.54±3.7 b | 9.48±1.38 a | 4.50±0.89 a |
| BIS C2 | 4.40±2.43 b | 0.25±0.14 c | 15.03±5.79 a | 7.51±1.70 b | 3.19±0.79 b |
| AMB C2 | 3.33±2.43 c | 0.45±0.27 a | 14.93±7.81 a | 7.24±1.81 b | 4.08±1.63 a |
| **Pr>F** | 2.2e-16*** | 2.2e-16*** | 3.505e-05*** | 1.473e-09*** | 2.466e-10*** |
| BOI C3 | 4.44±2.42 a | 0.22±0.07 b | 4.63±2.40 b | 9.21±1.52 a | 4.23±0.83 a |
| BIS C3 | 3.51±2 b | 0.16±0.09 c | 10.53±4.40 a | 7.19±1.52 b | 3.07±0.84 b |
| AMBC3 | 2.42±1.90 c | 0.31±0.10 a | 9.88±5.29 a | 6.64±1.80 b | 3.89±1.01 a |
| **Pr>F** | 2.2e-16*** | 2.2e-16*** | 1.86e-07*** | 1.319e-11*** | 1.847e-10*** |

**Table S4. Description and comparison of species based on total height and basal diameter variables.**

Total height and basal diameter vary according to the developmental stage of individuals across all species. Total height and basal diameter of the trunk differ significantly between species. At developmental stage 4, *C. bissetiae* shows the greatest height (358.73 ± 104.14 cm), while *C. boinensis* has the smallest height (190.61 ± 31.8 cm); *C. ambongensis* (249.73 ± 51.45 cm) is intermediate between the two. However, their basal diameters are relatively similar at this same physiological age. These species differ from one another at stages 1 and 2, particularly between *C. bissetiae* and *C. ambongensis*, in terms of basal diameter.

The following Table 12 provides the mean values (mean ± standard deviation); values sharing the same letters in a single row are not significantly different at the 5% significance level. Significance levels from the Kruskal-Wallis test are indicated as follows: * Significant, ** Highly significant, *** Very highly significant, and NS Not significant. AMB: *C. ambongensis*, BIS: *C. bissetiae*, BOI: *C. boinensis*.

| Development stages | Species | Total height (cm) | Basal diameter (cm) |
|---|---|---|---|
| 1 | BOI | 7.05±3.77 b | 0.13±0.02 ab |
| | BIS | 14.10±7.80 a | 0.21±0.05 a |
| | AMB | 14.63±5.26 ab | 0.22±0.08 b |
| | **Pr>F** | 0.039* | 0.0189 |
| 2 | BOI | 76.79±31.23 a | 0.79±0.18 ab |
| | BIS | 50.10±48.77 a | 0.48±0.33 b |
| | AMB | 109.51±28.4 a | 1.65±0.1 a |
| | **Pr>F** | 0.49 NS | 0.0208 ** |
| 3 | BOI | 189.99±106.55 a | 1.72±0.86 a |
| | BIS | 227.35±75.13 a | 1.56±0.43 a |
| | **Pr>F** | 0.62 NS | 0.708 NS |
| 4 | BOI | 190.61±31.8 b | 2.37±0.36 a |
| | BIS | 358.73±104.14 a | 3.20±0.79 a |
| | AMB | 249.73±51.45 ab | 3.96±1.5 a |
| | **Pr>F** | 0.0055** | 0.094 NS |

**Supplementary figures**

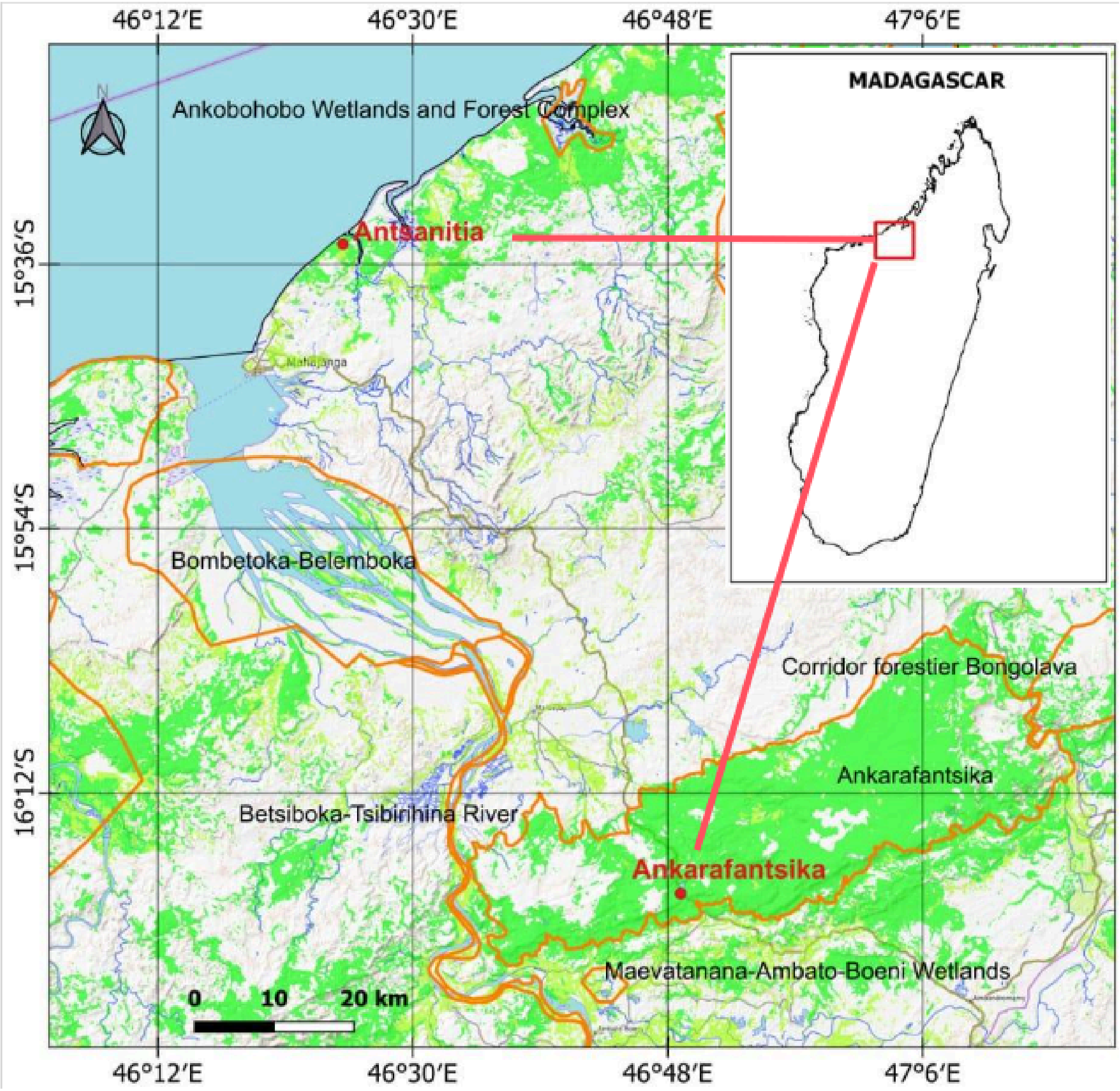

**Fig. S1. Location of the two main study sites.** Source: FTM Map (2016) modified by Bezandry (2023)

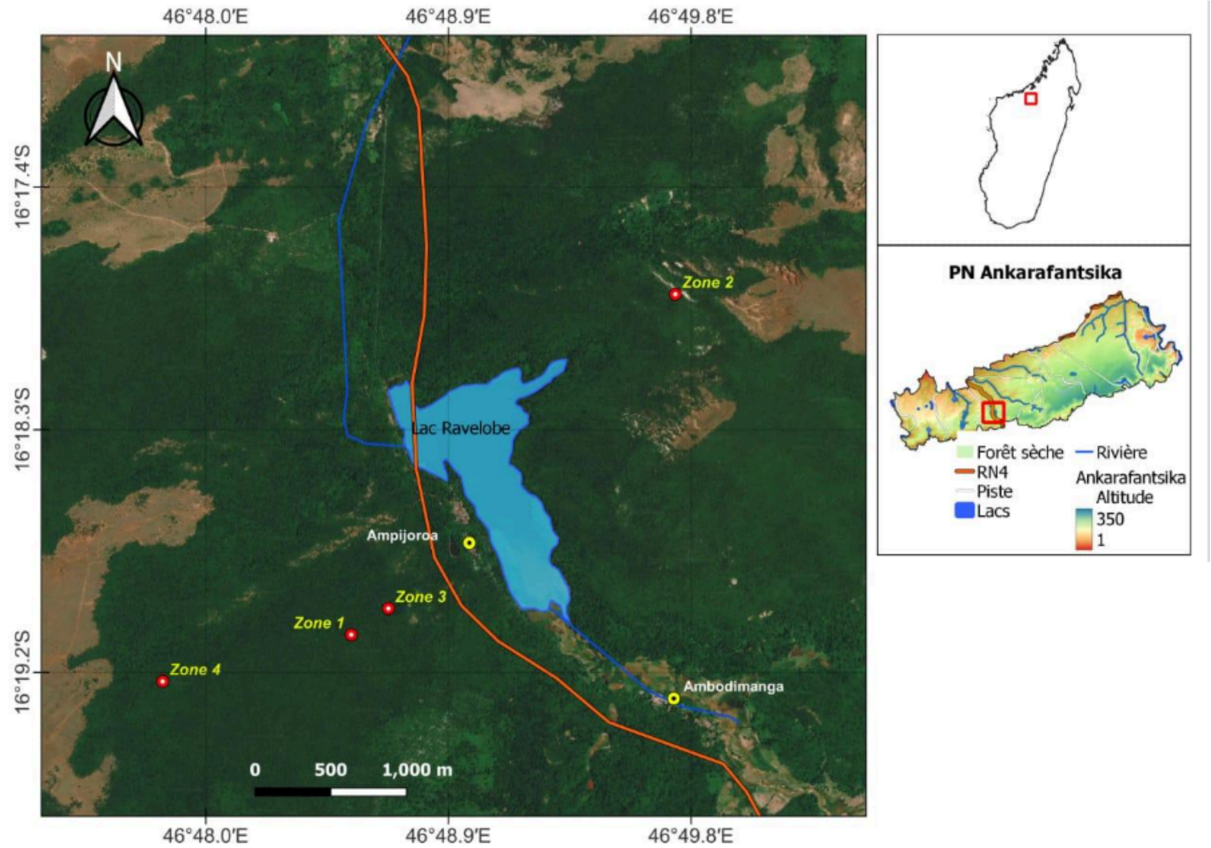

**Fig. S2. Location and subdivision of the study zones within PNA.** Source: FTM Map (2016) modified by Bezandry (2023).

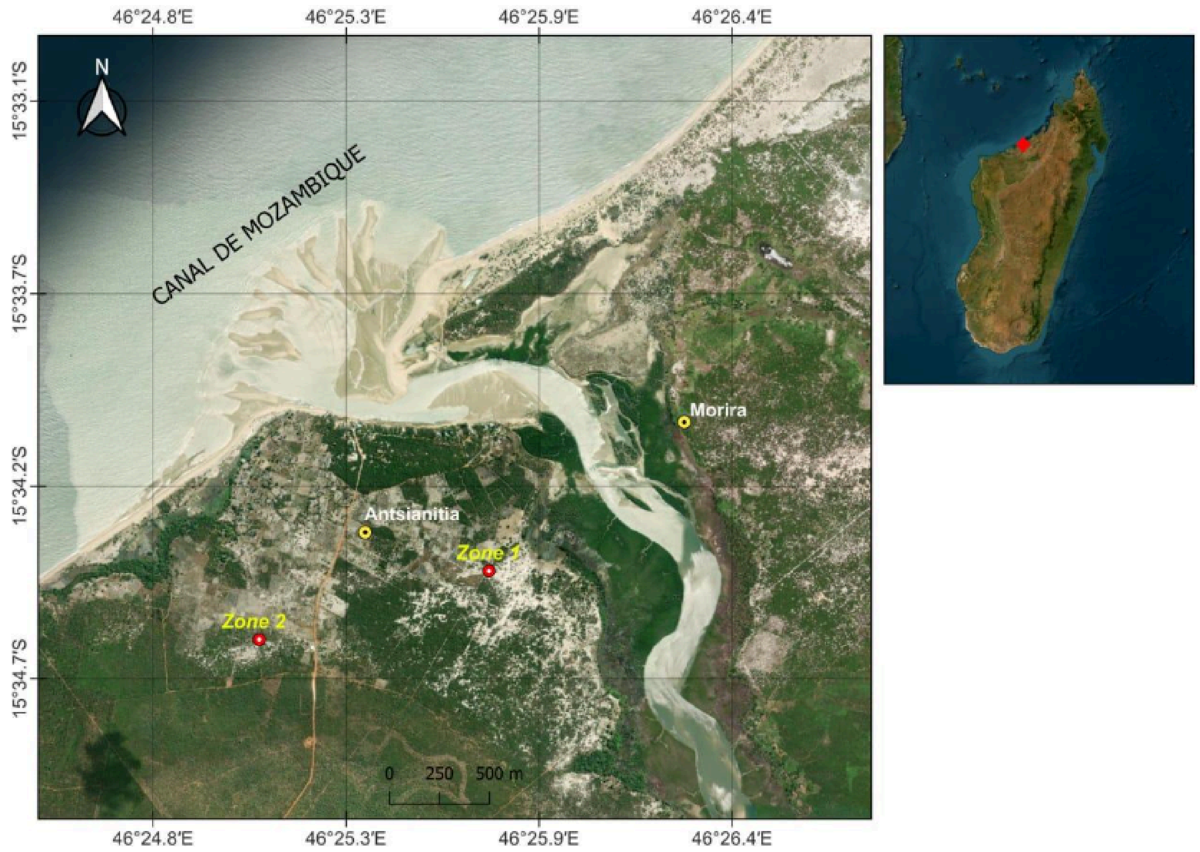

**Fig. S3. Location and subdivision of the study zones in Antsanitia.** Location and subdivision of the study zones in Antsanitia. Source: FTM Map (2016) modified by Bezandry (2023).

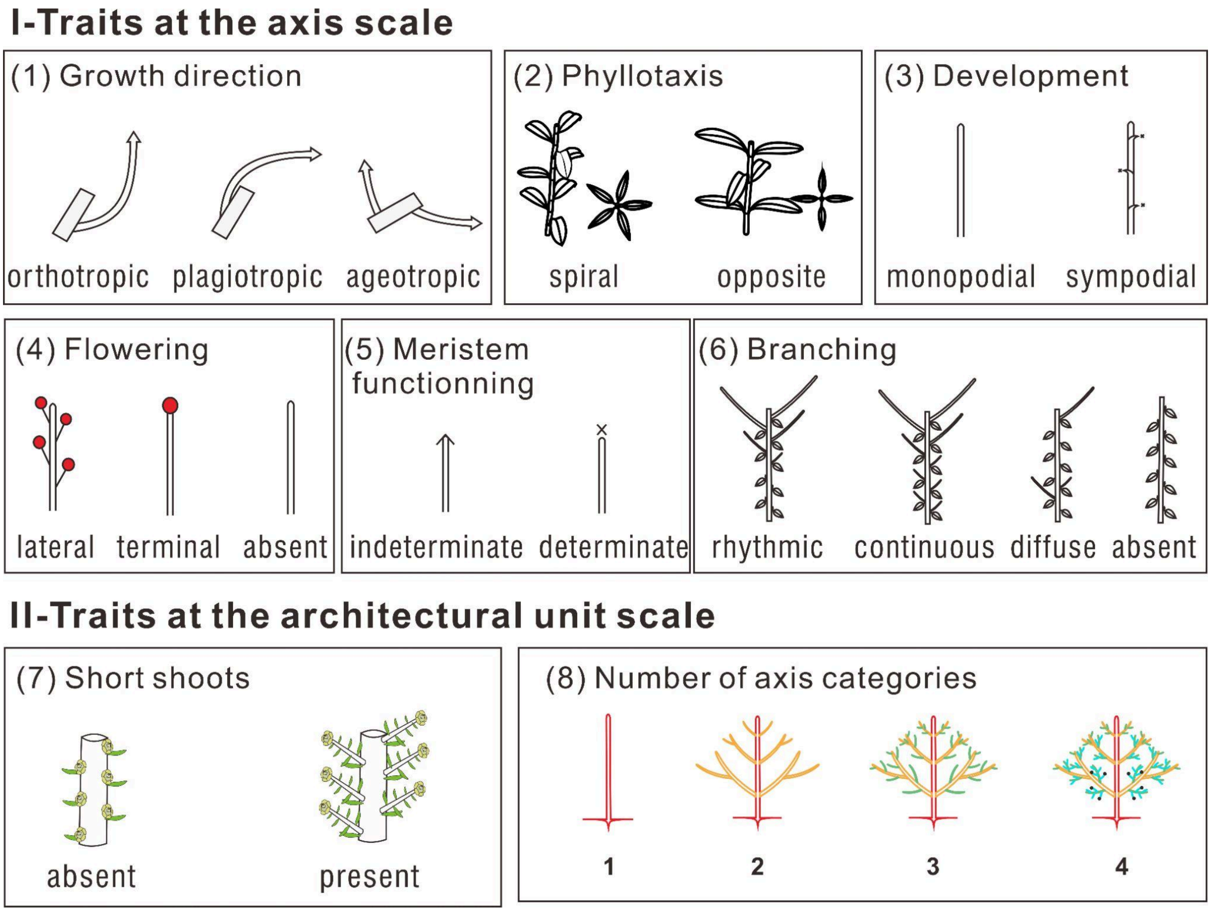

**Fig. S4. Some of the architectural descriptors commonly used in architectural analyses.** From Anest et al. (2021).

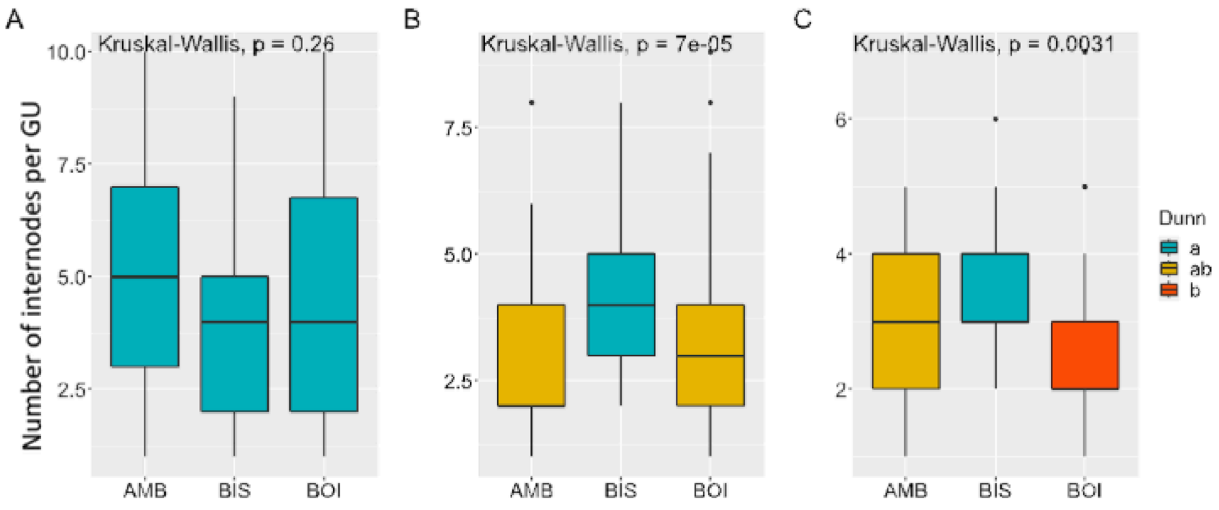

**Fig. S5. Statistical comparison (post-hoc) of the number of internodes per growth unit across species**

(A), (B), and (C) are graphical representations of the Kruskal-Wallis test results for the number of internodes per growth unit (GU) for the trunk, branches, and shoots, respectively. The Dunn test was used for pairwise comparisons (Post-Hoc). Species with the same colours are not statistically different at the 5% significance level.

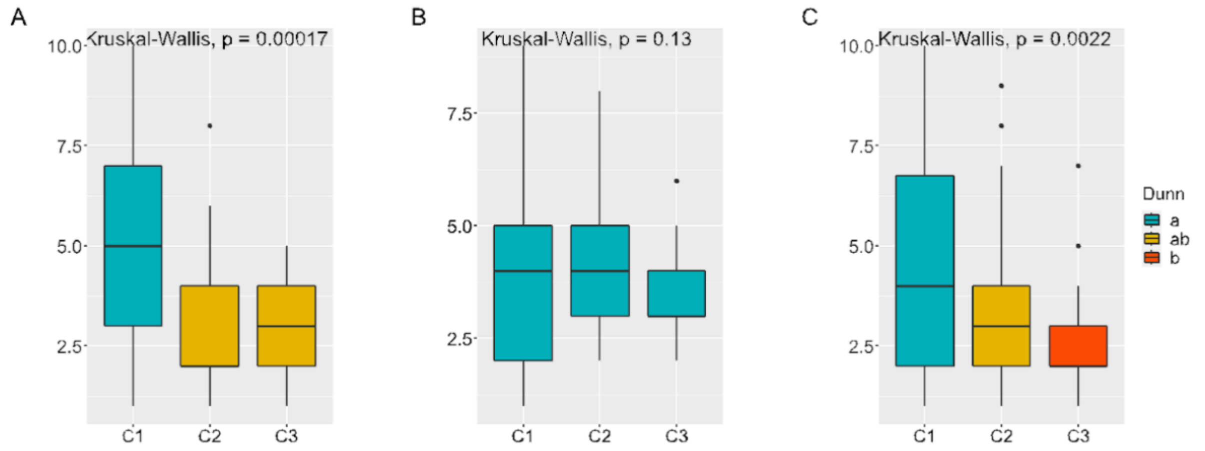

**Fig. S6. Statistical comparison (post-hoc) of the number of internodes per growth unit by axis type**

(A), (B), and (C) represent the Kruskal-Wallis test results for *C. ambongensis*, *C. bissetiae*, and *C. boinensis*, respectively. The axis types are trunks (C1), branches (C2), and shoots (C3). The Dunn test was used for pairwise comparisons (Post-Hoc). Axis types with the same colours are not statistically different at the 5% significance level.

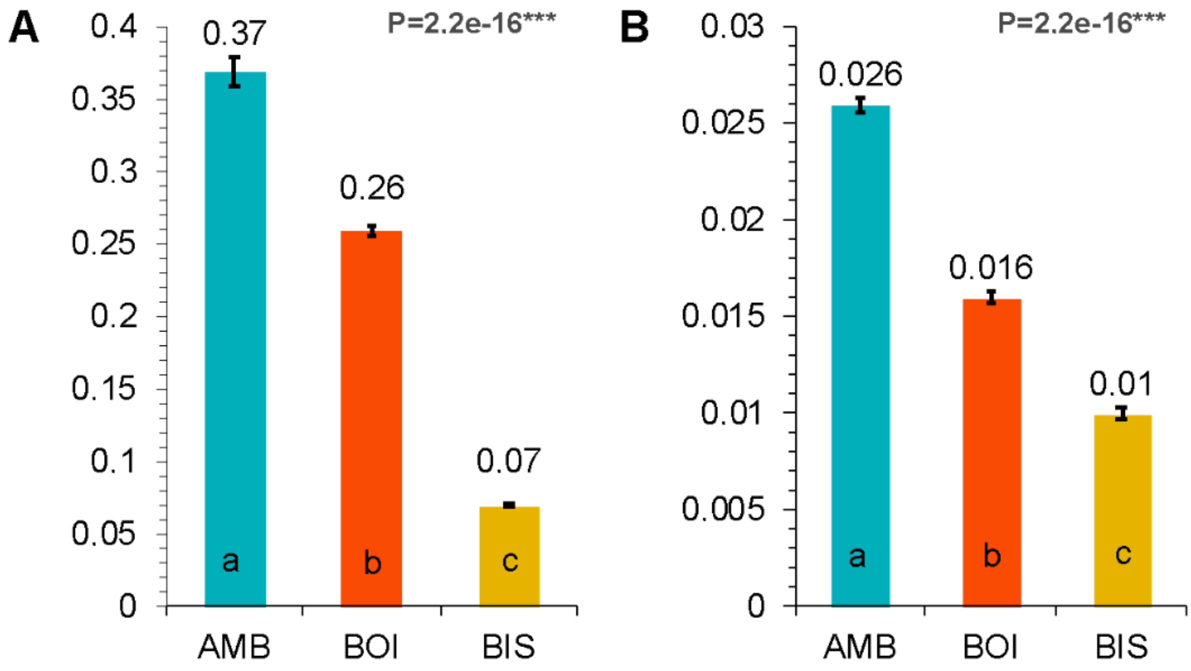

**Fig. S7. Average leaf dry mass and LMA by species**

(A) Average leaf dry mass (g) and (B) average leaf mass per area (LMA, g/cm²). Species are distinguished by colour codes. Species with the same letters (a, b, and c) are not statistically different at the 5% significance level.

Leaf dry mass (g) and LMA (g/cm²) vary significantly among the three species (p = 2.2e-16 ***). Leaf dry mass ranges from 0.07 g (*C. bissetiae*) to 0.37 g *(C. ambongensis*), while LMA ranges from 0.010 g/cm² (*C. bissetiae*) to 0.026 g/cm² (*C. ambongensis*). *C. ambongensis* has the highest leaf dry mass and LMA, while *C. bissetiae* has the lowest values.

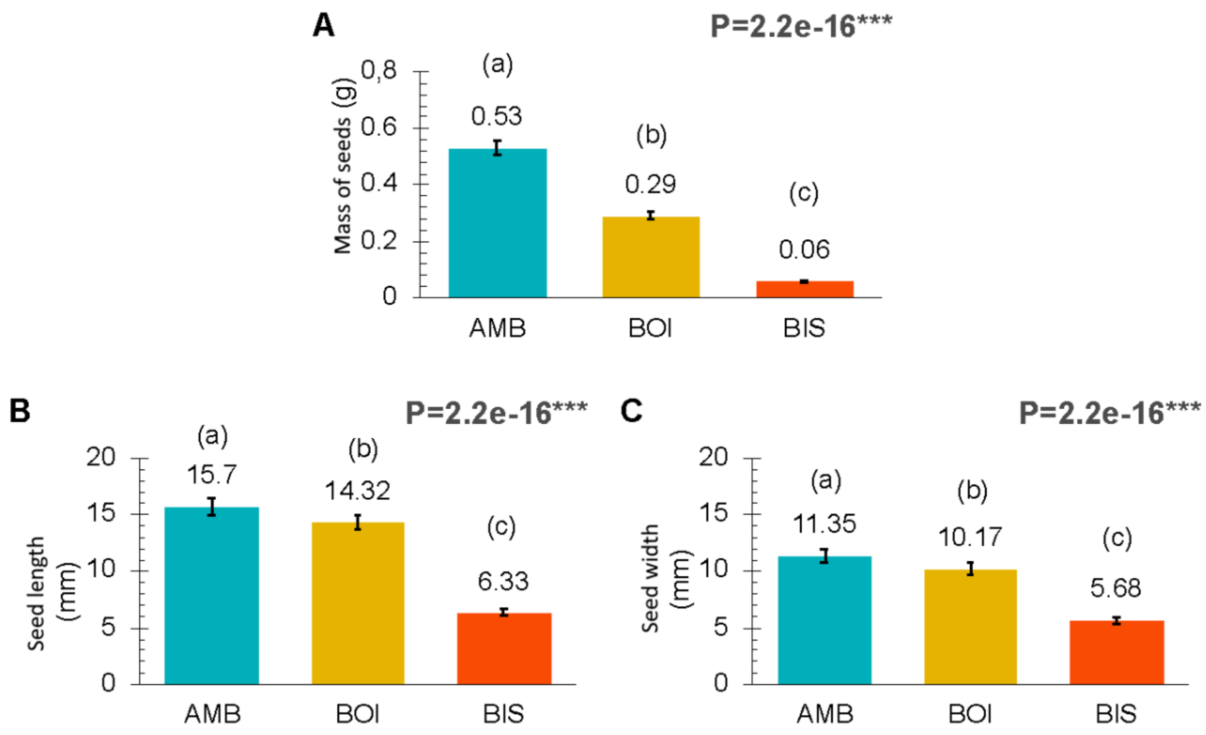

**Fig. S8. Average values of seed traits**

(A) Average seed mass (g), (B) seed length (mm), and (C) seed width (mm). Species are distinguished by colour codes. Species with the same letters (a, b, and c) are not statistically different at the 5% significance level.

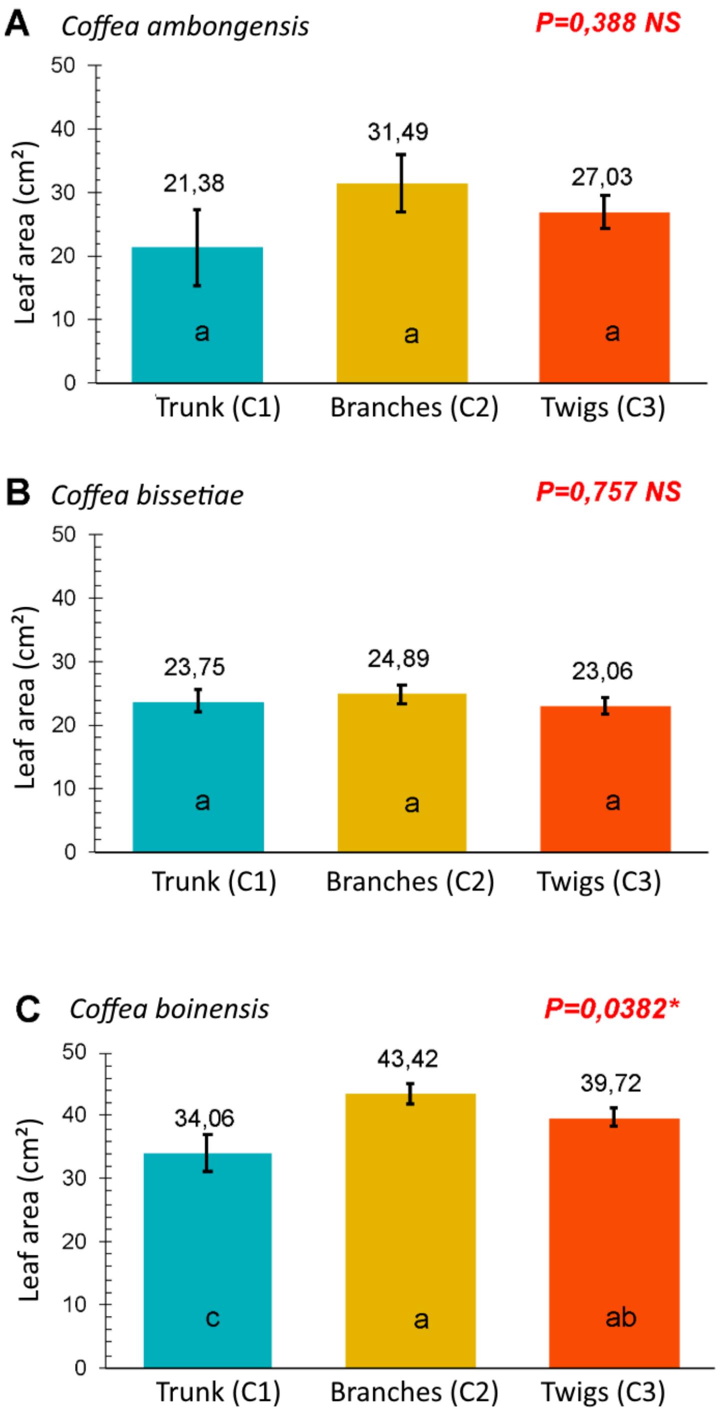

**Fig. S9. Average leaf area (length x width, in cm²) by axis category**

Leaf surface areas (length x width, in cm²) do not show significant differences across axis categories in individuals at stage 4 (A and B), except for branches and trunks in *C. boinensis* (P-value = 0.0382*; C). The difference is greater between branches and trunks than between branches and shoots, except in *C. bissetiae*. *C. boinensis* has the largest leaf surface area among the studied species.

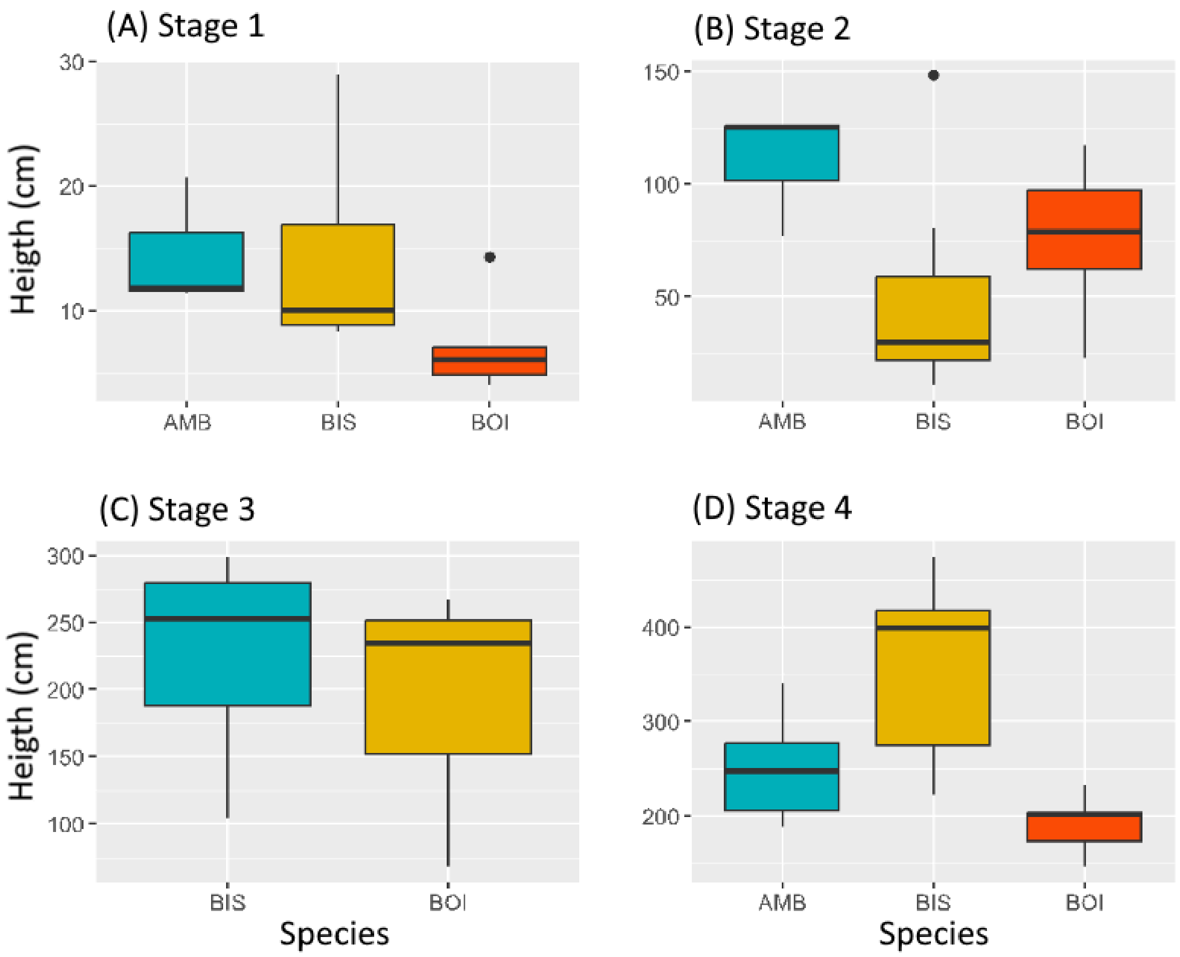

**Fig. S10. Height of the studied species across developmental stages**

(A), (B), (C), and (D) represent developmental stages (1), (2), (3), and (4), respectively. AMB: *C. ambongensis*, BIS: *C. bissetiae*, and BOI: *C. boinensis*.

At stage 1 of development, height ranges from 4 to 28.9 cm. On average, *C. ambongensis* is the tallest (14.62 ± 5.26 cm), *C. boinensis* is the shortest (7.05 ± 3.77 cm), and C. bissetiae has an intermediate height (14.1 ± 7.8 cm).

At stage 2, *C. ambongensis* is again the tallest, followed by *C. boinensis* and *C. bissetiae*, with heights ranging from 10.7 to 148 cm among the observed individuals.

At stage 3, heights range from 68.3 to 299.1 cm. *C. bissetiae* is the tallest, with an average height of 249.73 ± 51.45 cm . No individuals of *C. ambongensis* were observed at this developmental stage.

At stage 4, heights vary from 146.6 to 474.73 cm. *C. bissetiae* is the tallest (358 ± 104.14 cm), *C. boinensis* is the shortest (190.61 ± 31.8 cm), and *C. ambongensis* has an intermediate height (249.73 ± 51.45 cm).

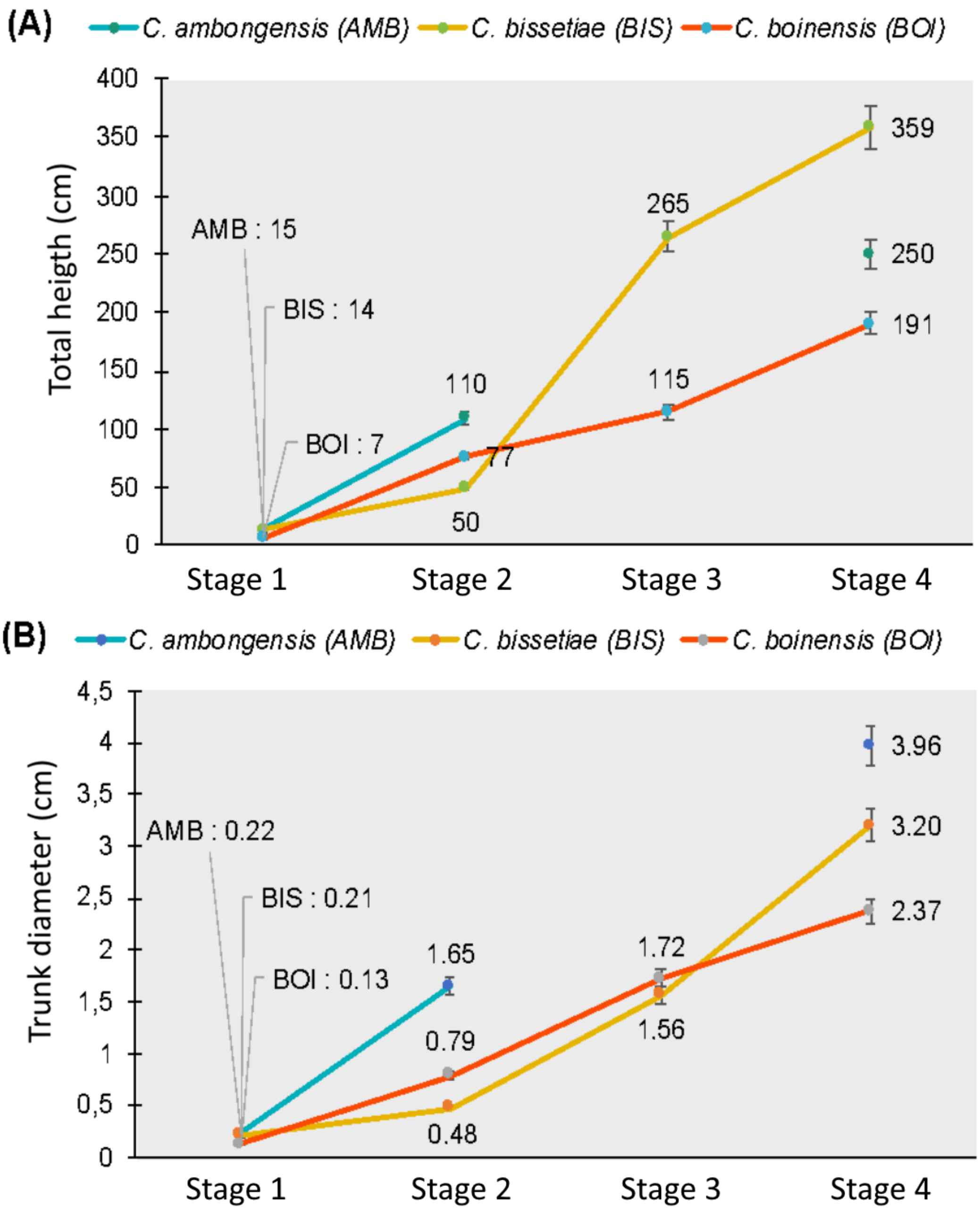

**Fig. S11. : Height and basal diameter from seedling to adult stage**

(A): Height and (B): Basal diameter of the observed individuals. At stage 1, height and diameter values are represented using grey lines and species initials to avoid overlapping values.The three species start with a more or less similar growth rhythm at stage 1 but differ significantly by stage 4. C. bissetiae, initially the shortest species at stage 1, rapidly increases in height starting from stage 2 and remains the tallest species at stages 3 and 4.

In terms of basal diameter, *C. bissetiae* follows a similar pattern but becomes the intermediate species at stage 4. In contrast, *C. boinensis* shows a more gradual increase in diameter throughout development.

A)

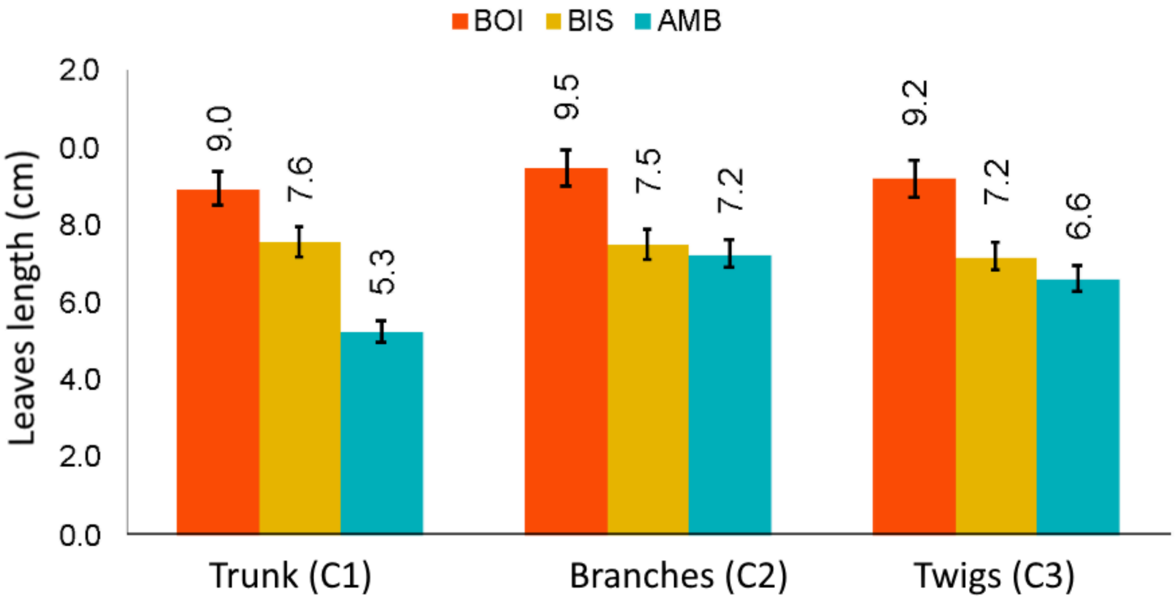

B)

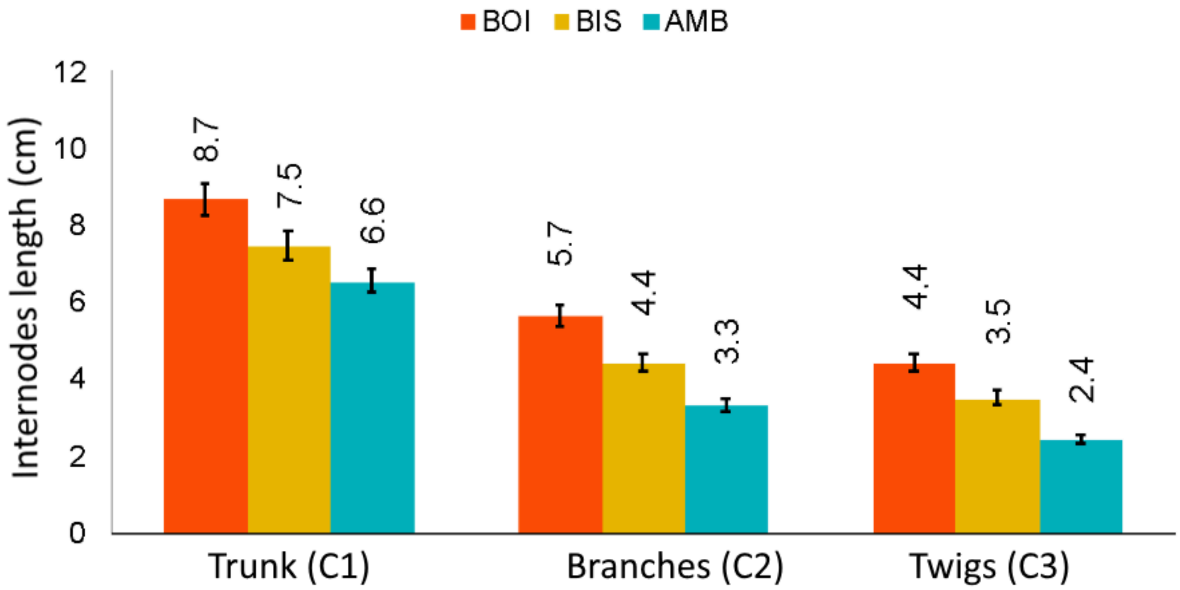

**Fig. S12. A) verage internode length and B) leaf length at developmental stage 4**

Species are distinguished by colour codes: BOI: *C. boinensis*, BIS: *C. bissetiae*, and AMB: *C. ambongensis.*

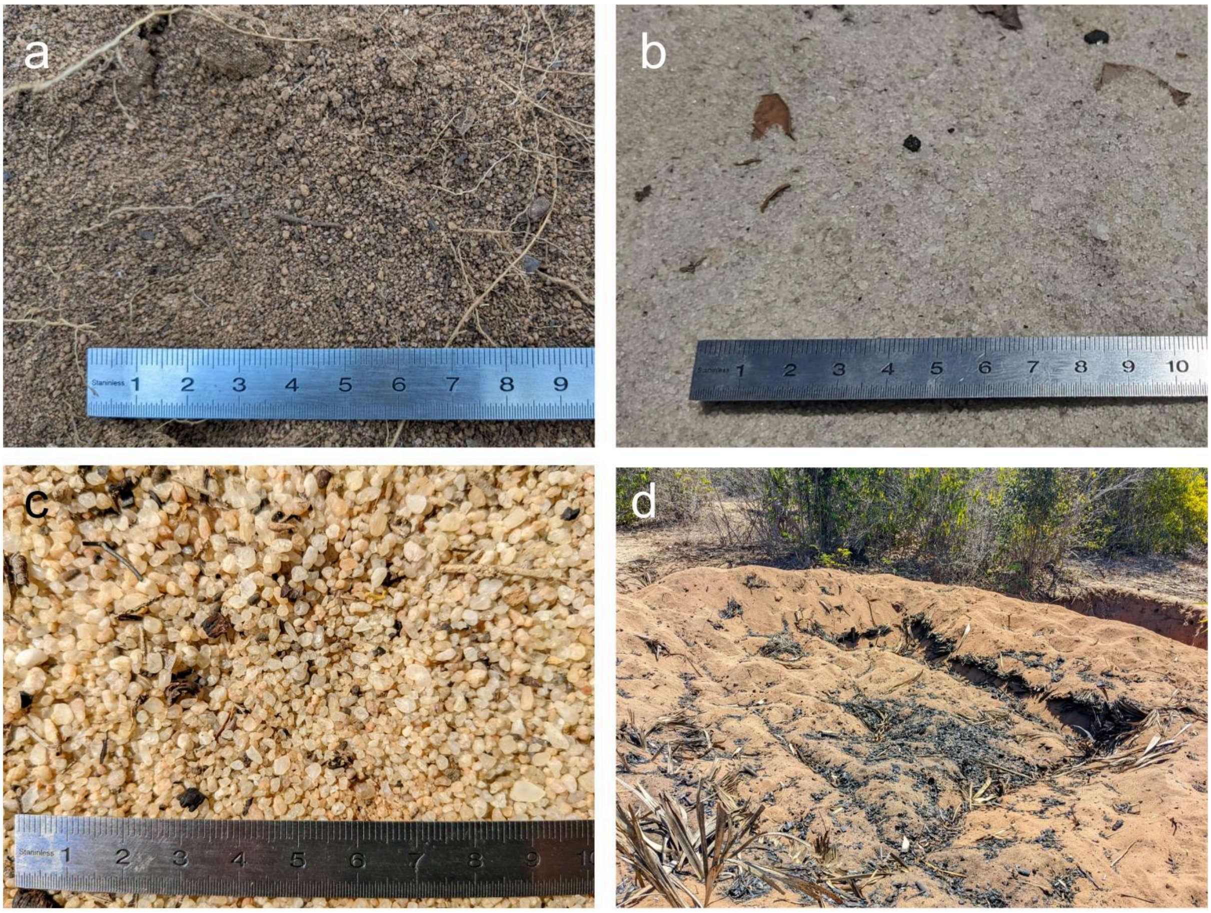

**Fig. S13. Substrate of the Baracoffea environments**

Substrate of a) *Coffea bissetiae* b) *Coffea ambongensis* c) *Coffea boinensis,* and d) rest of fire in *Coffea boninensis* environments.

**Notes. S1: Architectural and morphological description of Coffea ambongensis**

We identified four (4) categories of axes in the architectural unit of *Coffea ambongensis* (AMB), which differ in the traits described below.

The following table summarizes the traits of the four (4) categories of axes forming the architectural unit of the species C. ambongensis.

**Table S5: Summary of the architectural unit of *Coffea ambongensis* (AMB)**

| Descriptors | C1 | C2 | C3 | C4 |
|---|---|---|---|---|
| **Phyllotaxy** | Opposite-decussate | Opposite, secondary phyllotaxy | Opposite, secondary phyllotaxy | Alternate, rarely opposite |
| **Growth direction and symmetry** | Orthotropic | Plagiotropic | Plagiotropic | Plagiotropic |
| **Meristematic function and development** | Indeterminate, monopodial | Indeterminate, monopodial | Indeterminate, monopodial | Determinate, sympodial |
| **Growth rhythmicity** | Rhythmic | Rhythmic | Rhythmic | Rhythmic |
| **Branching in space** | Continuous | Continuous | Diffuse | Unbranched |
| **Branching in time** | Immediate | Immediate | Delayed | Unbranched |
| **Flowering** | Absent | Absent | Absent | Terminal |
| **Lifespan** | Plant lifetime | Very long | Very long | Short (3 GUs) |

**Trunk (C1)**

The trunk (C1) is an orthotropic monopod with opposite-decussate phyllotaxy (Fig. S14). The leaves are stipulate with a generally cordate blade shape (apex broadly acute to obtuse and margin smooth or slightly pubescent), although young leaves are occasionally oval. The leaves are slightly pubescent, and their size increases in shaded conditions. The blade is composed of a primary vein and secondary and tertiary veins.

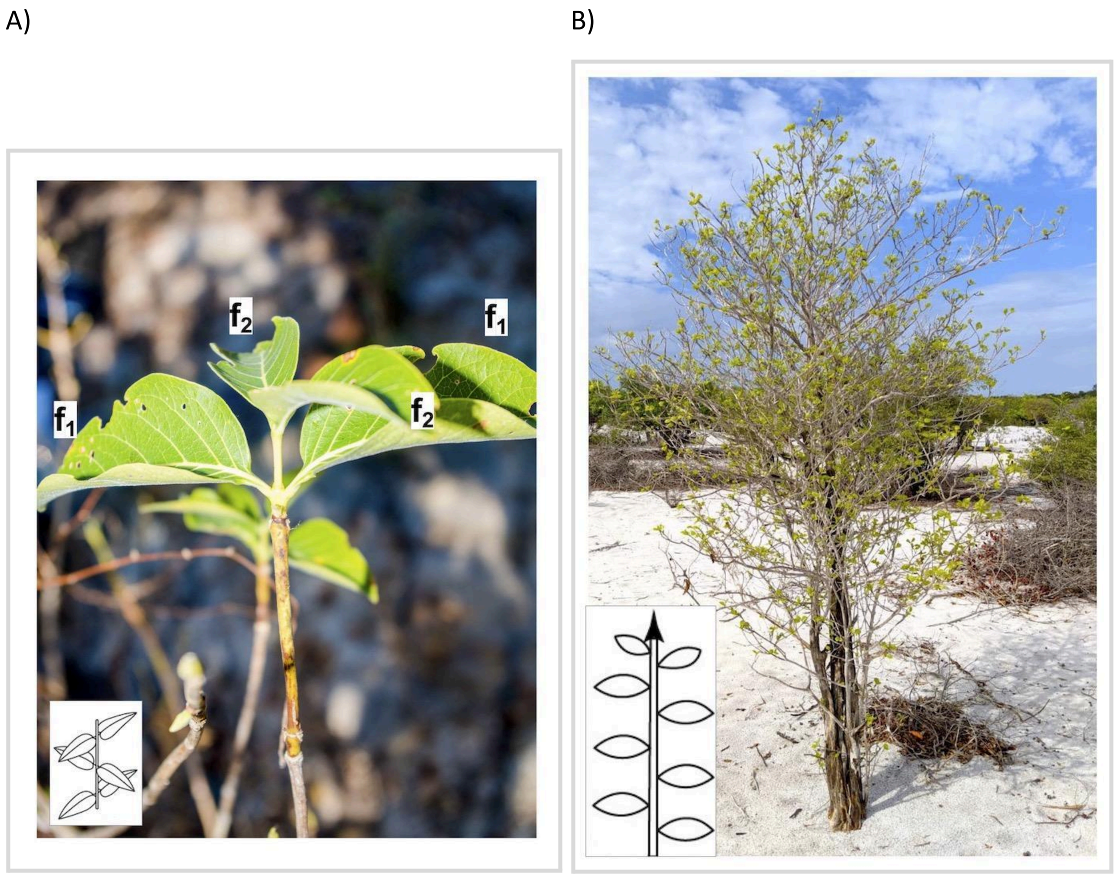

**Fig. S14. A) Opposite-decussate phyllotaxy in *C. ambongensis*; B) Branched system with monopodial development in *C. ambongensis*.** *f*: photosynthetic leaves.

The development of these axes (C1) is monopodial, characterized by an apical meristem that remains functional throughout the plant's lifetime, resulting in indeterminate growth (Fig. S14).

We observed a regular alternation between elongation phases and resting phases along the trunk. These phases are expressed as long internodes during elongation, with growth units delimited by closely spaced internodes marking growth slowdowns. Occasionally, reduced and scale-like leaves (cataphylls) are present at the growth arrest points. These traits indicate that the trunk has rhythmic growth (Fig. S15).

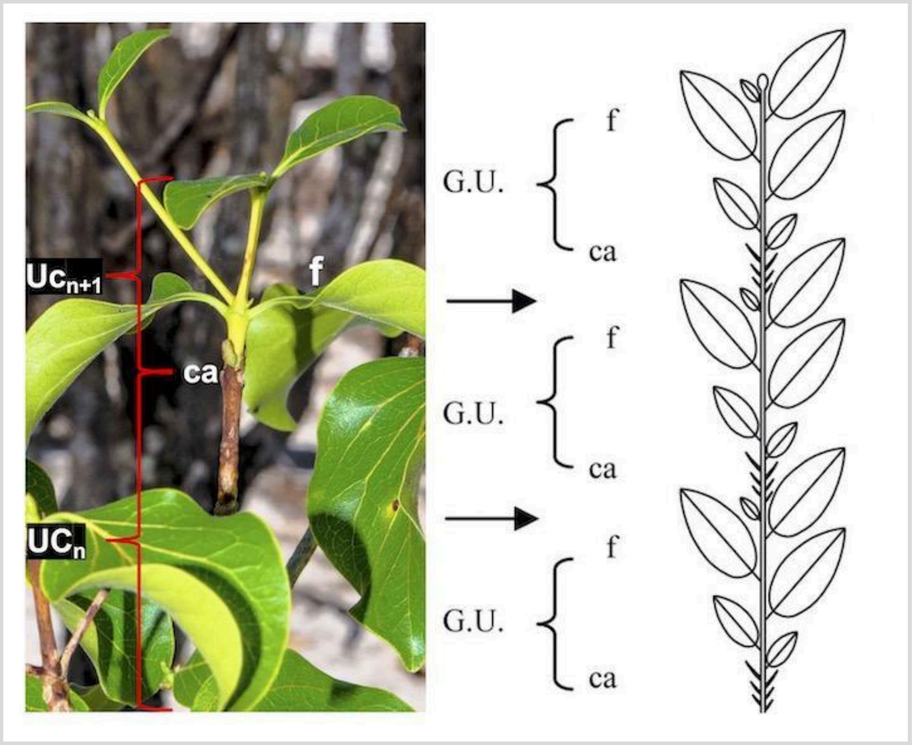

**Fig. S15: Rhythmic growth and morphological markers in *C. ambongensis*.** UC: Growth unit (a portion of the axis that elongated without interruption during an extension phase). ca: Cataphylls, which are morphological markers indicating growth stops. Cataphylls envelop and protect apical meristems during resting phases in unfavorable periods. Older growth units appear brown, while newer ones are light green.

The trunk's branching is usually continuous, showing a basitonic gradient (towards the base of the growth units), though it can occasionally be diffuse. Branching is often immediate (sylleptic) but can also occasionally be delayed (proleptic).

Immediate branching typically begins at the first node formed (often a short node) in the new growth unit following a growth slowdown. At this point, two axes frequently develop in the axils of the opposite leaves (Fig. S16A). Less commonly, only one of the two buds develops (Fig. S16C).

In contrast, delayed branching often begins in the axil of a long node (and strictly not at the first node formed in a new growth unit) (Fig. S16B). Delayed branching leading to plagiotropic axes is less frequent at the base.

The trunk (C1) is one of the axis categories that persist throughout the plant's life. It does not bear reproductive structures.

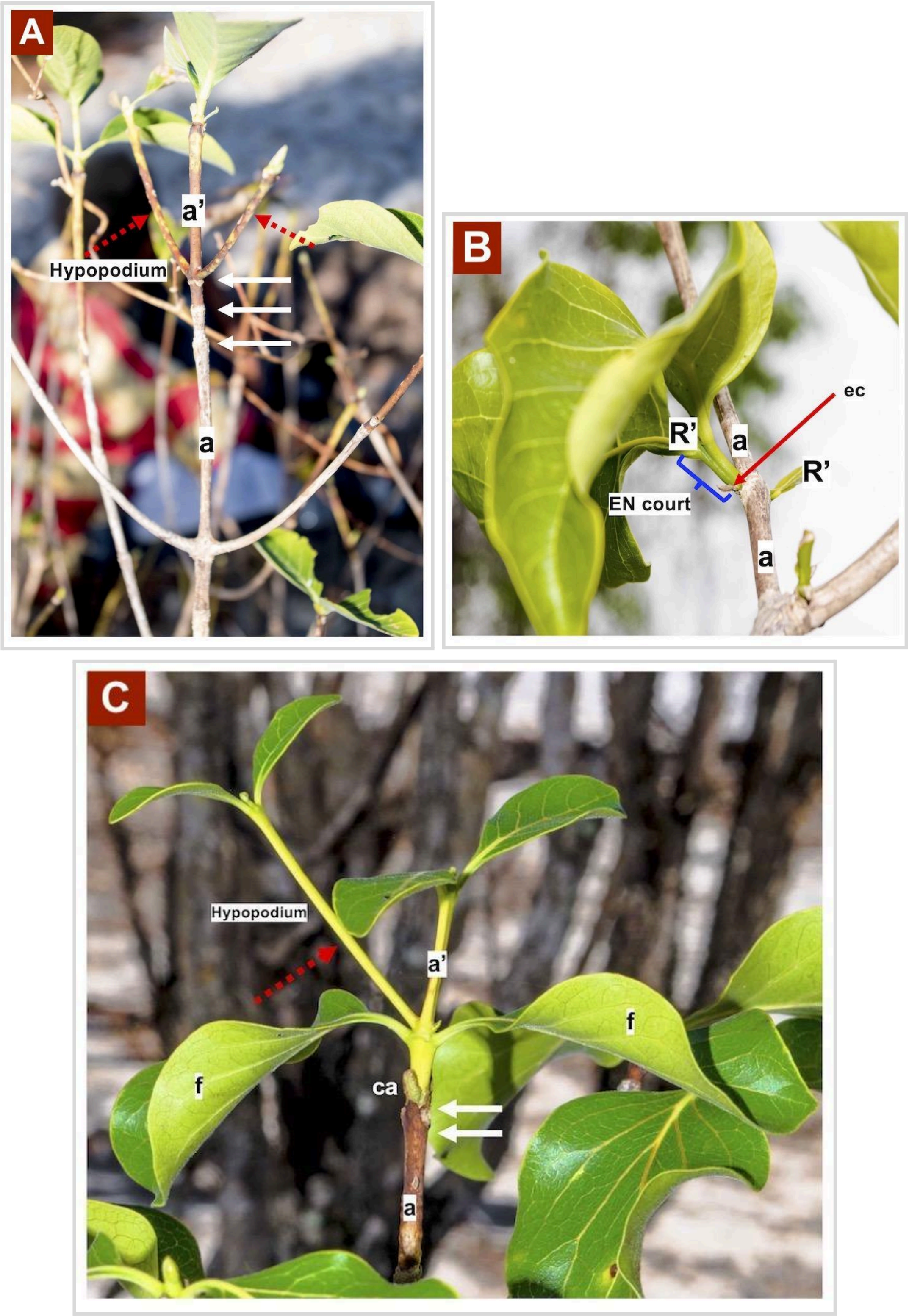

**Fig. S16: Immediate and delayed branching in *C. ambongensis*** A) Immediate branching on growing axes. B) Delayed branching on axes that have completed elongation. C) Immediate branching at the first node formed in the new growth unit, with a single branch inserted in the axil of two opposite leaves. a: Old shoot; a': New shoot; ca: Cataphylls; f: Assimilatory leaves; ec: Scales; R': Newly

branched axes on the fully elongated axes (with significantly smaller size and a different color compared to the supporting axis).
White arrows indicate growth arrests (translated by closely spaced internodes), and red dashed arrows indicate branches resulting from immediate branching (i.e., branches whose supporting axes are still elongating), which are characterized by longer initial internodes (hypopodium).
For delayed branching, during the elongation phase of the supporting axes (C1), the axillary buds enter dormancy after their formation and are protected by scales or cataphylls (ec). These dormant buds develop only after a certain period. When these dormant buds elongate, the first internodes formed are relatively short (short EN).

**Branches (C2):**
Branches (C2) are monopodial plagiotropic axes with opposite phyllotaxy, where leaves are secondarily arranged in a single plane through petiole twisting. They show monopodial development with an indeterminate meristematic function. Growth interruptions are marked by closely spaced internodes, indicating rhythmic growth. Their branching is lateral and predominantly continuous (all buds develop). Axes that develop laterally on branches can originate from newly formed buds on the elongating growth unit (immediate branching) or from latent buds (delayed branching). While immediate branching produces twigs, delayed branching (frequently observed on branches) gives rise to axes morphologically similar to the parent branch. Branches (C2) have a very long lifespan, equivalent to that of the plant. No sexuality has been observed on branches.

**Twigs (C3):**
Twigs (C3) are monopodial plagiotropic axes with opposite phyllotaxy. Similar to branches, their leaves are secondarily arranged in a single plane through petiole twisting. These twigs show monopodial development with an indeterminate meristematic functioning, although it can rarely become determinate. In rare cases of determinate function, we observed apical meristems transforming into inflorescences after several phases of initially indeterminate meristematic activity. Their growth is rhythmic, with growth interruptions marked by closely spaced internodes. Like the trunk (C1) and branches (C2), twigs (C3) persist throughout the plant's lifespan. Twigs (C3) show lateral branching, predominantly delayed. They rarely bear reproductive organs (terminally positioned when present).

**Short Shoots (C4):**
Short shoots (C4) are sympodial plagiotropic axes. They initially show alternate phyllotaxy during

vegetative growth, transitioning to opposite phyllotaxy when bearing reproductive structures. These axes have sympodial development with determinate meristematic function. Their growth is rhythmic, characterized by very short internodes. Short shoots branch sympodially (mono- or dichasially). Sexuality is generally terminal (transformation of the apical meristem into a flower) and rarely lateral (Fig. S17). After fruit maturation, one (occasionally two) successors develop sympodially from lateral buds located just below the terminal flowering zone.

The lifespan of short shoots (C4) is significantly shorter than that of other axis categories (C1, C2, and C3). They can be pruned after several flowering/fruiting seasons.

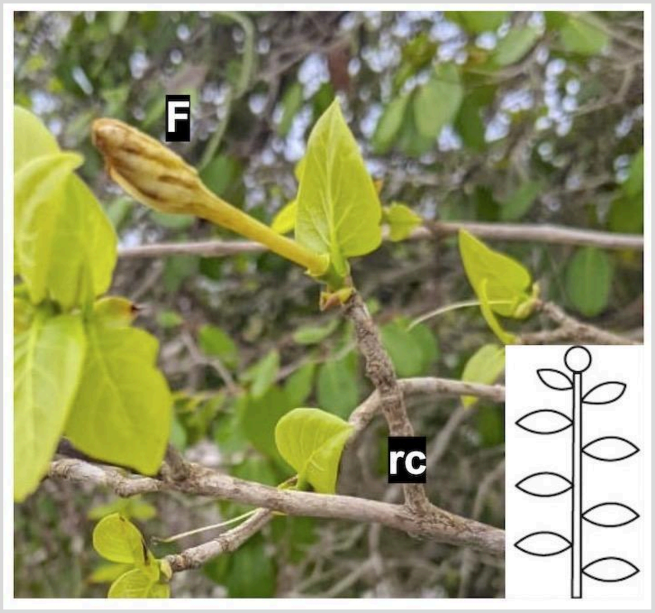

**Fig. S17: Terminal Sexuality on Short Shoots in *Coffea ambongensi*s**. F: flower and rc: short shoots**.** The "rc" axes are the only axis category capable of developing flowers from their apical meristems, indicating that they are the only axes with sympodial development.

**Notes. S2: Architectural and morphological description of *Coffea bissetiae***

We identified four (4) axis categories in the architectural unit of C. bissetiae, which differ based on the traits described below.

**Table S6: Summary of the Architectural Unit of *Coffea bissetiae***

| Descriptors | C1 | C2 | C3 | C4 |
|---|---|---|---|---|
| **Phyllotaxy** | Opposite-decussate | Opposite, secondary phyllotaxy | Opposite, secondary phyllotaxy | Alternate, rarely opposite |
| **Growth direction and symmetry** | Orthotropic | Plagiotropic | Plagiotropic | Plagiotropic |
| **Meristematic function and development** | Indeterminate, monopodial | Indeterminate, monopodial | Indeterminate, monopodial | Determinate, sympodial |
| **Growth rhythmicity** | Rhythmic | Rhythmic | Rhythmic | Rhythmic |
| **Branching in space** | Continuous | Continuous | Diffuse | Unbranched |
| **Branching in time** | Immediate | Immediate | Delayed | Unbranched |
| **Flowering** | Absent | Absent | Absent | Terminal |
| **Lifespan** | Plant lifespan | Very long | Very long | Short (3 growth units) |

**Trunk (C1)**

The main axis (C1) has opposite-decussate phyllotaxy. It is a monopodial orthotropic axis that can vary in length depending on environmental conditions: it remains relatively short in the open environment of Antsanitia, whereas it can grow very tall in the forested, closed environment of Ankarafantsika (Fig. S18).

The leaves are simple (moderately pubescent), stipulate, and petiolate, with an elliptical (sometimes lanceolate or oval) lamina that is slightly pubescent. The lamina comprises a primary vein, secondary veins, and tertiary veins. The leaf morphology is consistent across different axis categories.

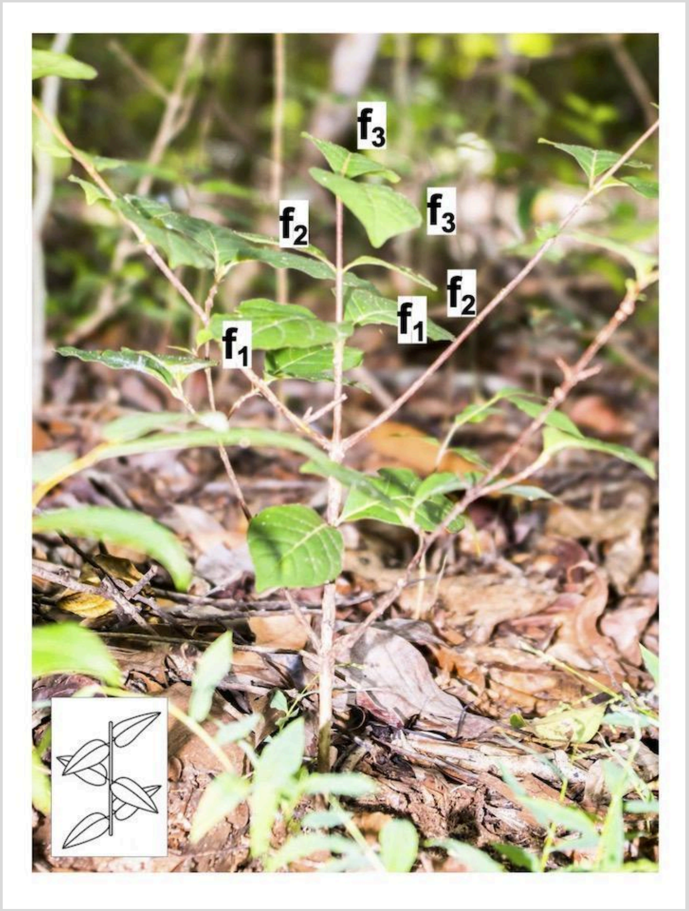

**Fig. S18: Opposite-Decussate Phyllotaxy in *Coffea bissetiae*.** f: assimilative leaves

The trunk (C1) shows an indeterminate meristematic function (Fig. S19). A regular alternation of long and short internodes was observed, indicative of rhythmic growth. Growth pauses are marked by closely spaced internodes, the presence of protective transformed leaves (cataphylls) safeguarding the apical meristem during the dormant phase, and a color change in the axis (Fig. S19).

The trunk (C1) remains intact throughout the plant's lifespan. Branching is predominantly continuous and immediate (Fig. S20). The trunk does not bear reproductive structures.

A) B)

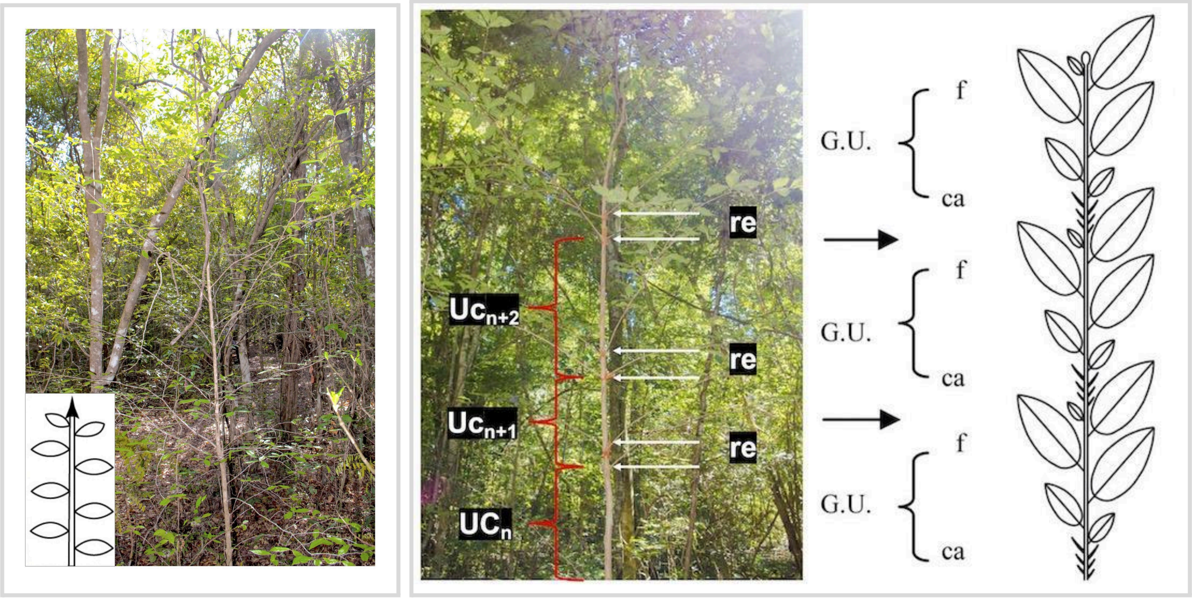

**Fig. S19. A) Monopodial branched system in *Coffea bissetiae*; B) Rhythmic growth and morphological markers in *Coffea bissetiae*.** UC: Growth unit (portions of axes that elongate without interruption during an extension phase). re: Closely spaced internodes (EN), which are morphological markers indicating growth pauses.

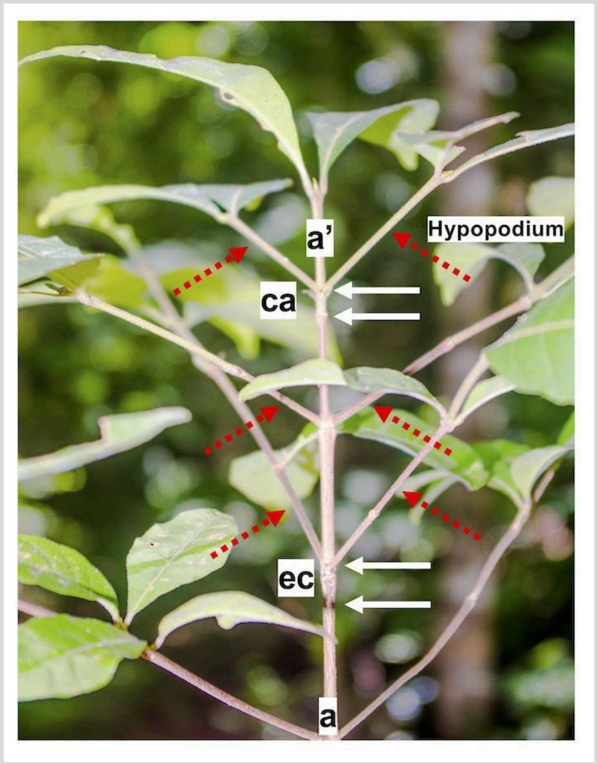

**Fig. S20. Immediate branching in *Coffea bissetiae*.** a: Old shoot; a': New shoot; ca: Cataphylls; f: Assimilative leaves; ec: Scales.

White arrows indicate growth pauses (represented by closely spaced internodes, EN), and red dashed arrows indicate branches resulting from immediate branching (i.e., branches originating from portions of the trunk still elongating), characterized by long initial internodes (hypopodia). All nodes show branching (continuous branching).

**Branches (C2)**

Branches (C2) are monopodial, plagiotropic axes with opposite, non-decussate phyllotaxy. Leaves are secondarily arranged in a single plane due to petiole torsion.

The branches have an indefinite meristematic functioning and rhythmic growth. Growth interruptions are often marked by closely spaced internodes, the presence of reduced leaves at the base of a growth unit, and changes in bark coloration preceding the interruption. Newly formed growth units are lighter in color, whereas older growth units are darker. Branches (C2) persist throughout the plant's lifespan. Branch branching is continuous and typically immediate.

**Twigs (C3)**

Twigs (C3) are monopodial, plagiotropic axes with opposite phyllotaxy. Leaves are secondarily arranged in a single plane due to petiole torsion.

Twigs have indefinite meristematic functioning and rhythmic growth. Similar to the trunk (C1) and branches (C2), twigs (C3) persist throughout the plant's lifespan. Twigs have diffuse and delayed branching. They do not bear reproductive structures.

**Short shoots (C4)**

Short shoots (C4) are sympodial, plagiotropic axes. These are relatively short axes, typically bearing a single leaf (rarely two) at the base of their growth.

These axes (C4) show definite meristematic functioning and rhythmic growth. The apical meristem of short shoots has seasonal floral activity, with a relay axis (rarely two) developing through delayed branching after fruit maturation or abortion. Growth is rhythmic, alternating between an elongation phase and a resting phase. Resting phases are characterized by the formation of cataphylls (or scales) of a brownish color and very short internodes.

The previous season's growth unit is gray to brown, while the new growth unit is slightly flattened, pubescent, and light green-brown.

Reproduction is strictly terminal, involving the transformation of the apical meristem into a floral bud (Fig. S21). Unlike trunks (C1), branches (C2), and twigs (C3), short shoots (C4) have a short lifespan, typically consisting of 2-3 growth units before being pruned.
Occasionally, short shoots have been observed developing apically (after a growth interruption) on branches (C2).

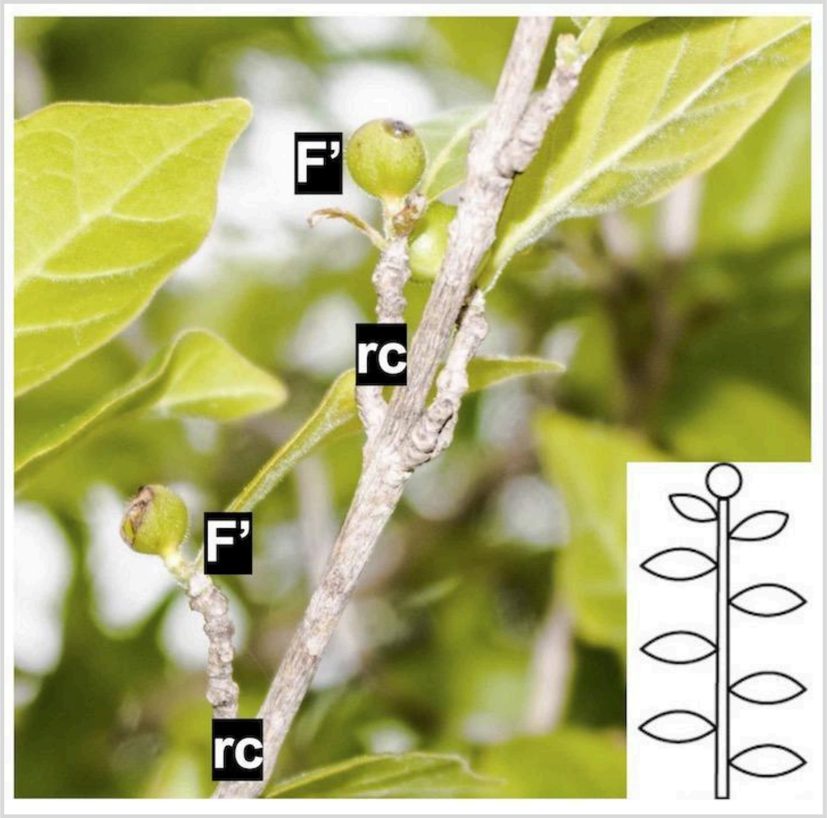

**Fig. S21: Terminal sexuality on short shoots in *Coffea bissetiae*** F': Maturing fruit; rc: Short shoots. On this figure, the rc are borne by twigs (C3).

**Notes S3: Architectural and morphological description of Coffea boinensis**

We identified four (4) categories of axes in the architectural unit of Coffea boinensis, which differ in the traits described below.

**Table S7. Recap of the architectural unit of Coffea boinensis (BOI)**

| Descriptors | C1 | C2 | C3 | C4 |
|---|---|---|---|---|
| **Phyllotaxy** | Opposite-decussate | Opposite, secondary plane | Opposite, secondary plane | Opposite, rarely alternate |
| **Growth direction and symmetry** | Orthotropic | Plagiotropic | Plagiotropic | Plagiotropic |
| **Meristematic function and development** | Indeterminate, monopodial | Indeterminate, monopodial | Indeterminate, monopodial | Determinate, sympodial |
| **Growth rhythm** | Rhythmic | Rhythmic | Rhythmic | Rhythmic |
| **Branching in space** | Continuous | Continuous | Diffuse | Unbranched |
| **Branching in time** | Immediate | Immediate | Delayed | Unbranched |
| **Flowering** | Absent | Absent | Absent | Terminal |
| **Lifespan** | Entire plant life | Very long | Very long | Very short (2 UC) |

**Trunk (C1)**

The trunk (C1) is a monopodial orthotropic axis with opposite-decussate phyllotaxy (Fig. S22). Leaves are simple, stipulate, and petiolate, with a lamina that is deltoid, cordate, or oval in shape, with a pointed apex and an entire margin. The lamina is composed of a primary vein, secondary veins, and tertiary veins.

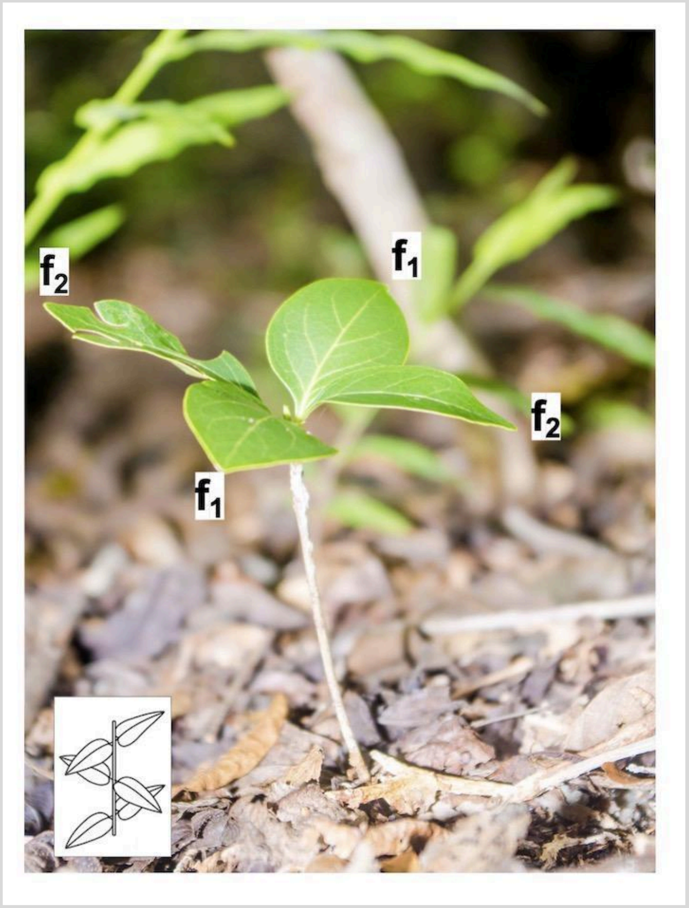

**Fig. S22: Opposite-decussate phyllotaxy in *Coffea boinensis*.** f: Photosynthetic leaves.

The trunk has an indeterminate meristematic function and rhythmic growth (Fig. S23). Growth occurs seasonally, producing a new growth unit (UC), which is greenish-brown and flattened, composed of one or more long internodes. Older growth units are grey or brown in color. Growth stops are often marked by short internodes and the presence of cataphylls (Fig. S23A).

The trunk (C1) has the same lifespan as the plant. Its branching is continuous and immediate (Figs. S23B, S24). The trunk does not bear any reproductive structures.

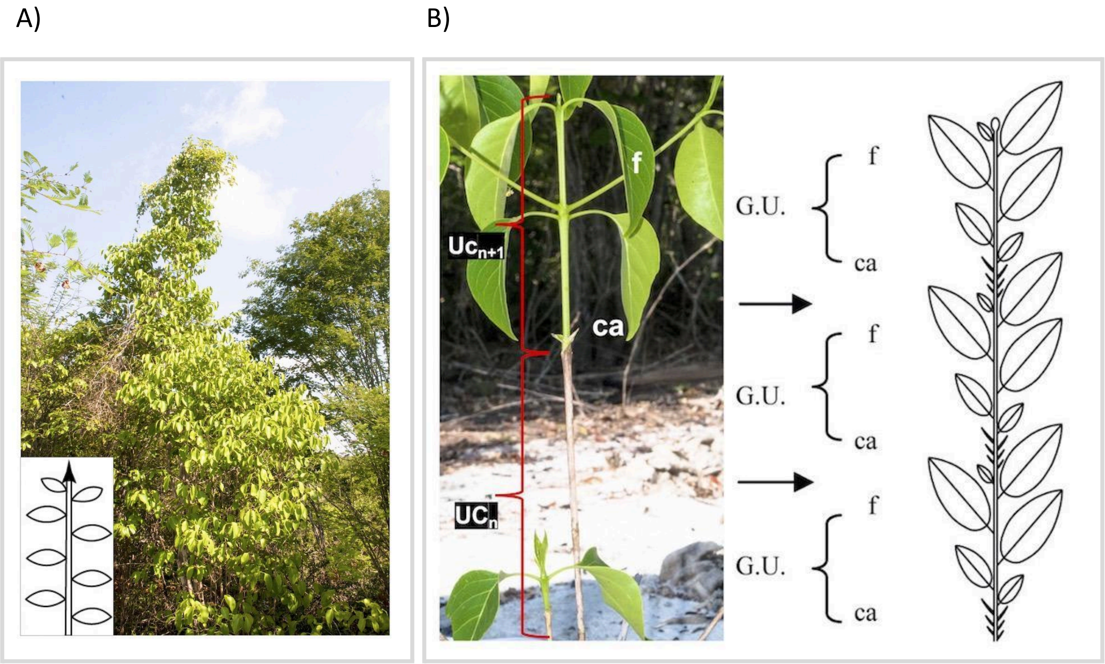

**Fig. S23 : A) Monopodial branching system in *Coffea boinensis*. B) Rhythmic growth and morphological markers in *Coffea boinensis.*** UC: Growth unit (these are portions of axes that elongated continuously during an extension phase); ca: Cataphylls, which are morphological markers indicating growth stops. The cataphylls surround and protect the apical meristems during dormancy phases in unfavorable periods. Older growth units are grey in color, while new growth units are light green.

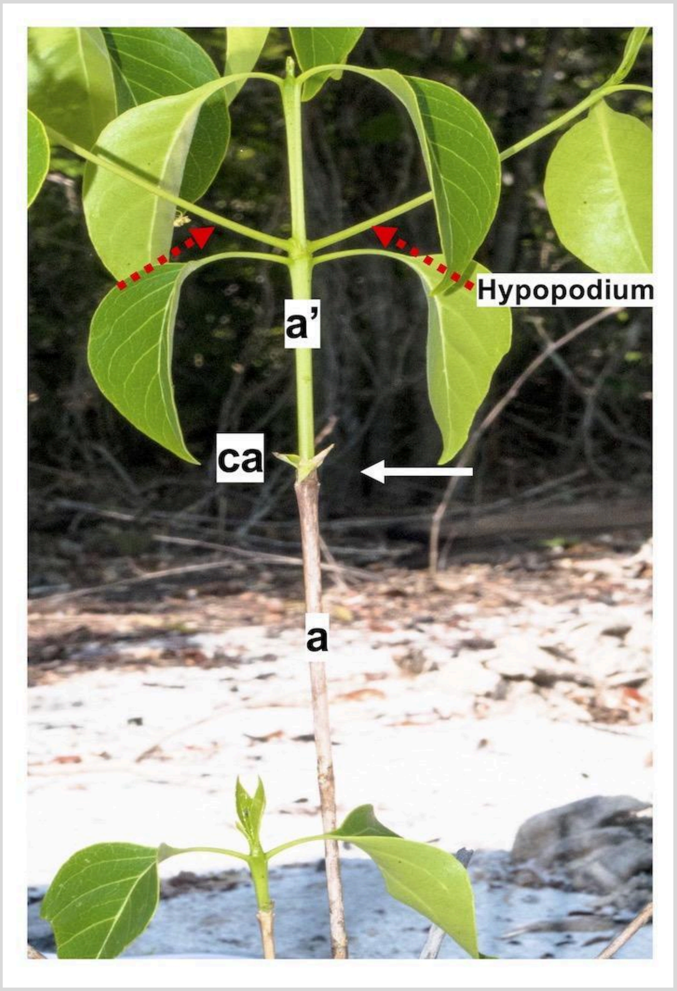

**Figure S24. Immediate branching in *Coffea boinensis*.** a: Axis that has completed its elongation; a': Axis currently elongating; ca: Cataphylls separating two successive growth units, a and a'.

The dashed red arrows indicate branches resulting from immediate branching (i.e., development of branches (C2) on a portion of the trunk (C1) still elongating). These new branches consist of long initial internodes (hypopodia).

Branches (C2)

Branches (C2) are monopodial plagiotropic axes with opposite phyllotaxy. Leaves are secondarily arranged in a single plane through petiole torsion. They show indefinite, rhythmic growth, characterized by seasonal flushes composed of one or more long internodes following a dormancy phase. Growth markers, such as short internodes and protective cataphylls during dormancy, are identical to those observed on the trunk.

Branches are long-lived, persisting throughout the plant's lifespan. Their branching is predominantly immediate and continuous (though delayed branching can occasionally be observed in older axes). Newly formed and branched axes differ in color (green-brown and flattened) and diameter (smaller than the supporting axis, which is grey or brown and larger). Branches (C2) do not bear sexual organs.

**Twigs (C3)**

Twigs (C3) are monopodial plagiotropic axes with opposite phyllotaxy and leaves arranged in a single plane due to petiole torsion. Twigs show indefinite rhythmic growth and persist throughout the plant's lifespan. Their branching is generally delayed (occasionally immediate) and diffuse.

Twigs do not bear reproductive structures.

**Short Shoots (C4)**

Short shoots (C4) are sympodial plagiotropic axes. These axes are relatively short, with a total cumulative length of only a few centimeters. Typically, they bear a single leaf (rarely two), located at the base of the terminal floral organ or fruit.

Short shoots show a defined meristematic function and rhythmic growth. Their apical meristem is involved in floral development, and a sympodial relay axis may develop (from a dormant lateral bud located just below the fruit) after the fruit matures or aborts (sometimes).

Short shoot growth is characterized by very small internodes marking growth stops and very short growth units compared to other axis categories (C1, C2, and C3). The short shoot is distinguished by an initial relatively long growth unit at the base, followed by shortened subsequent internodes. Growth stops are marked by very short, scaly, swollen, brown, or grey internodes. Growth units are characterized by long internodes on new growth with a green-brown color.

These axes are the only ones to bear reproductive structures in a terminal position (Fig. S25).

Short shoots have a relatively short lifespan (2-3 growth units before being pruned).

Short shoots frequently develop on branches (C2).

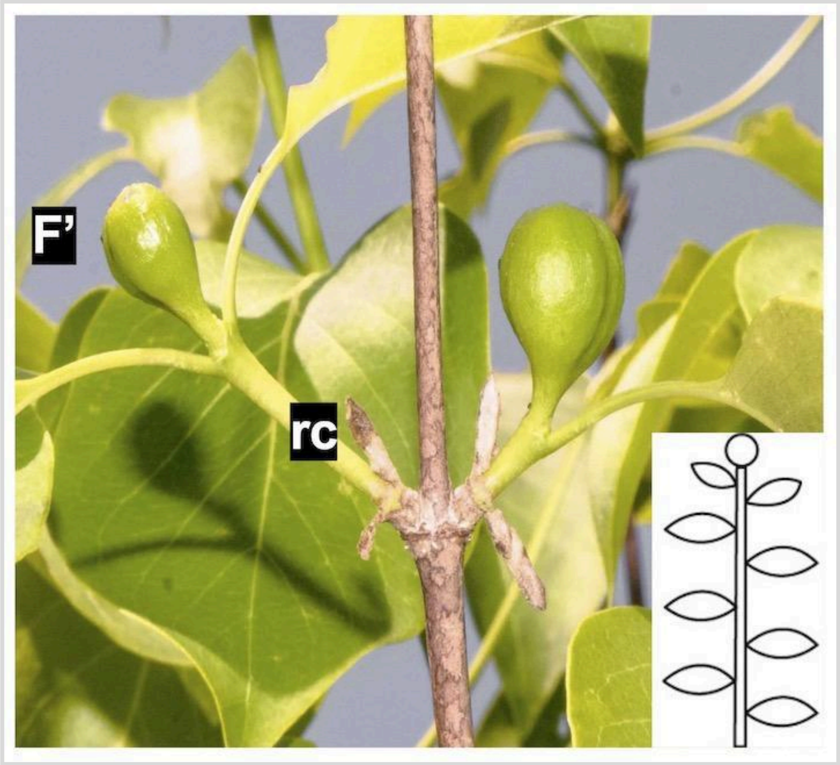

**Fig. S25: Terminal sexuality in *Coffea boinensis***; F’: Maturing fruits; rc: Short shoots.

In this figure, the short shoots (rc) are borne on branches (C2). The development of the short shoots occurred through delayed branching.
